# *Gpr158* deficiency impacts hippocampal CA1 neuronal excitability, dendritic architecture, and affects spatial learning

**DOI:** 10.1101/506295

**Authors:** Demirhan Çetereisi, Ioannis Kramvis, Titia Gebuis, Rolinka J. van der Loo, Yvonne Gouwenberg, Huibert D. Mansvelder, Ka Wan Li, August B Smit, Sabine Spijker

**Author notes:** Equal contribution (Demirhan Çetereisi, Ioannis Kramvis).

## Abstract

G-protein-coupled receptor 158 (*Gpr158*) is highly expressed in striatum, hippocampus and prefrontal cortex. It gained attention as it was implicated in physiological responses to stress and depression. Recently, *Gpr158* has been shown to act as a pathway-specific synaptic organizer in the hippocampus, required for proper mossy fiber-CA3 neurocircuitry establishment, structure, and function. Although rodent *Gpr158* expression is highest in CA3, considerable expression occurs in CA1 especially after the first postnatal month. Here, we combined hippocampal-dependent behavioral paradigms with subsequent electrophysiological and morphological analyses from the same group of mice to assess the effects of *Gpr158* deficiency on CA1 physiology and function. We demonstrate deficits in spatial memory acquisition and retrieval in the Morris water maze paradigm, along with deficits in the acquisition of extinction memory in the passive avoidance test in *Gpr158* KO mice. Electrophysiological recordings from CA1 pyramidal neurons revealed normal basal excitatory and inhibitory synaptic transmission, however, Schaffer collateral stimulation yielded dramatically reduced post-synaptic currents. Interestingly, intrinsic excitability of CA1 pyramidals was found increased, potentially acting as a compensatory mechanism to the reductions in Schaffer collateral-mediated drive. Both *ex vivo* and *in vitro,* neurons deficient for *Gpr158* exhibited robust reductions in dendritic architecture and complexity, i.e., reduced length, surface, bifurcations, and branching. This effect was localized in the apical but not basal compartment of adult CA1 pyramidals, indicative of pathway-specific alterations. A significant positive correlation between spatial memory acquisition and extent of complexity of CA1 pyramidals was found. Taken together, we provide first evidence of significant disruptions in hippocampal CA1 neuronal dendritic architecture and physiology, driven by *Gpr158* deficiency. Importantly, the hippocampal neuronal morphology deficits appear to support the impairments in spatial memory acquisition observed in *Gpr158* KO mice.

## Introduction

G-protein-coupled receptors (GPCRs) form a large family of seven transmembrane proteins. In the brain, these act as important regulators of synaptic transmission, neuronal excitability, and structural plasticity, through presynaptic and postsynaptic mechanisms of action (Huang and Thathiah, 2015; Leung and Wong, 2017). Several brain-expressed GPCRs were shown to modulate cognition and are suspect in the pathobiology of neuronal disorders (Catapano and Manji, 2007; Thompson et al., 2008). Collectively, GPCRs make up the largest drug target group to-date (Chan et al., 2015). Recently, G-protein coupled receptor 158 (*Gpr158)* caught attention due to its role in hippocampus-dependent memory formation via osteocalcin (OCN) signaling (Khrimian et al., 2017). *Gpr158* is found highly expressed in the brain, specifically in prefrontal regions, striatum, and hippocampus (Khrimian et al., 2017; Orlandi et al., 2012; Sutton et al., 2018). Previously, we have observed Gpr158 strongly upregulated in the hippocampus synaptic membrane fraction after stress of an immediate shock (Rao-Ruiz et al., 2015). The stress-responsiveness of *Gpr158* was further illustrated by its rapid and robust increase in expression upon glucocorticoid stimulation either *in vitro* (Patel et al., 2013), or *in vivo* (Sutton et al., 2018). Finally, *Gpr158* was shown increased in the prefrontal cortex (PFC) in the hours to days after physical stress in mice, as well as in post-mortem prefrontal tissue from depressed patients, whereas *Gpr158* KO mice exhibited resilience to unpredicted chronic mild stress (Sutton et al., 2018).

Recent work has begun shedding light on the precise function of *Gpr158* (Condomitti et al., 2018; Khrimian et al., 2017; Orlandi et al., 2012; Sutton et al., 2018). *Gpr158* overexpression in mouse medial PFC (mPFC) increases helplessness, illustrated by increased immobility in a tail-suspension (TST) and forced-swim tests (FST) (Sutton et al., 2018). On the contrary, global deletion of *Gpr158* had an anxiolytic and anti-depressant effect, exhibited by increased number of crossovers and times spent in the open arm of an elevated plus maze (EPM) (Sutton et al., 2018). Prefrontal reestablishment of *Gpr158* expression in *Gpr158* KO mice equalized these EPM differences. Recordings from *Gpr158* KO mPFC superficial layer pyramidals revealed increases in spontaneous excitatory postsynaptic current (sEPSC) frequency, and an AMPA receptor (AMPAR) current-driven increase in the AMPAR/NMDAR ratio, partially supported by an increase in spine density (Sutton et al., 2018).

Hippocampal-dependent learning appears negatively affected in *Gpr158* KO mice, as observed during Morris water maze (MWM) training, and as shown by memory deficits in the novel object recognition task (NOR) (Khrimian et al., 2017). Dentate gyrus’ (DG) mossy fiber (MF) to CA3 long-term potentiation (LTP) was shown significantly impaired in these mice (Khrimian et al., 2017). Moreover, frequency and amplitude of sEPSCs, AMPAR and NMDAR evoked currents, and MF to CA3 paired pulse ratio (PPR) were all robustly reduced in *Gpr158* KO CA3 pyramidals (Condomitti et al., 2018). Although CA3 spine density and MF-CA3 synapse density were paradoxically found increased, both the area and volume per MF-CA3 synapse were robustly reduced (Condomitti et al., 2018). Importantly, the aforementioned electrophysiological and morphological deficits were pathway-specific, affecting MF to CA3 synapses, but not CA3 to CA3 association/commissural synapses. The underlying mechanism for this was elegantly demonstrated by the discovery of glypican 4 (Gpc4), which is highly enriched in MFs, as the presynaptic binding partner to postsynaptic hippocampal *Gpr158* (Condomitti et al., 2018).

Given the considerable CA1 expression from post-weaning onward (Condomitti et al., 2018), as well as the role of *Gpr158* in neuronal signaling and cognition (Khrimian et al., 2017; Sutton et al., 2018), we investigated the effect of *Gpr158* deficiency on both electrophysiological properties and neuronal architecture in the hippocampus CA1 with *in vitro* and *ex vivo* approaches. We took specific advantage of measuring hippocampus-dependent behavior and performing electrophysiological recordings with cellular reconstructions in the same set of animals, allowing us to create insightful correlations between the behavior and underlying neuronal architecture observed. Taken together, our data reveal spatial learning deficits and neuronal hyperexcitability in *Gpr158* KO mice, driven by the compromised dendritic architecture of CA1 pyramidal neurons.

## Results

### Gpr158 *KO mice show aberrant spatial learning and decreased acquisition of safety learning*

Given the significant hippocampal expression of *Gpr158* (Allen brain atlas probe RP_051121_01_A10; (Khrimian et al., 2017), we tested *Gpr158* KO mice and WT littermates in an array of hippocampal-dependent behavioral paradigms (Figure 1 supplement 1). First, we performed MWM training with male Bl6/J *Gpr158* KO mice (**Figure 1a,b**, Figure 1 supplement 1) and a probe test to assess long-term memory retainment of hidden platform location based on distal cues. During the 4-day training protocol, the distance swam to find the hidden platform was found statistically different in *Gpr158* KO mice *vs*. WT mice (**Figure 1a**, mixed ANOVA: genotype *P*=0.046; training *P*=0.500, interaction *P*=0.059), with post-hoc testing showing increased distance in *Gpr158* KO mice *vs*. WT mice during the last 2 training days (t-test: day3 *P*=0.031, day 4 *P*=0.025). In addition, escape latency showed a similar effect (Figure 1 supplement 1 and supplement 2), albeit the first training day showed an initial genotype difference in activity (Figure 1 supplement 1).

**Figure 1.**
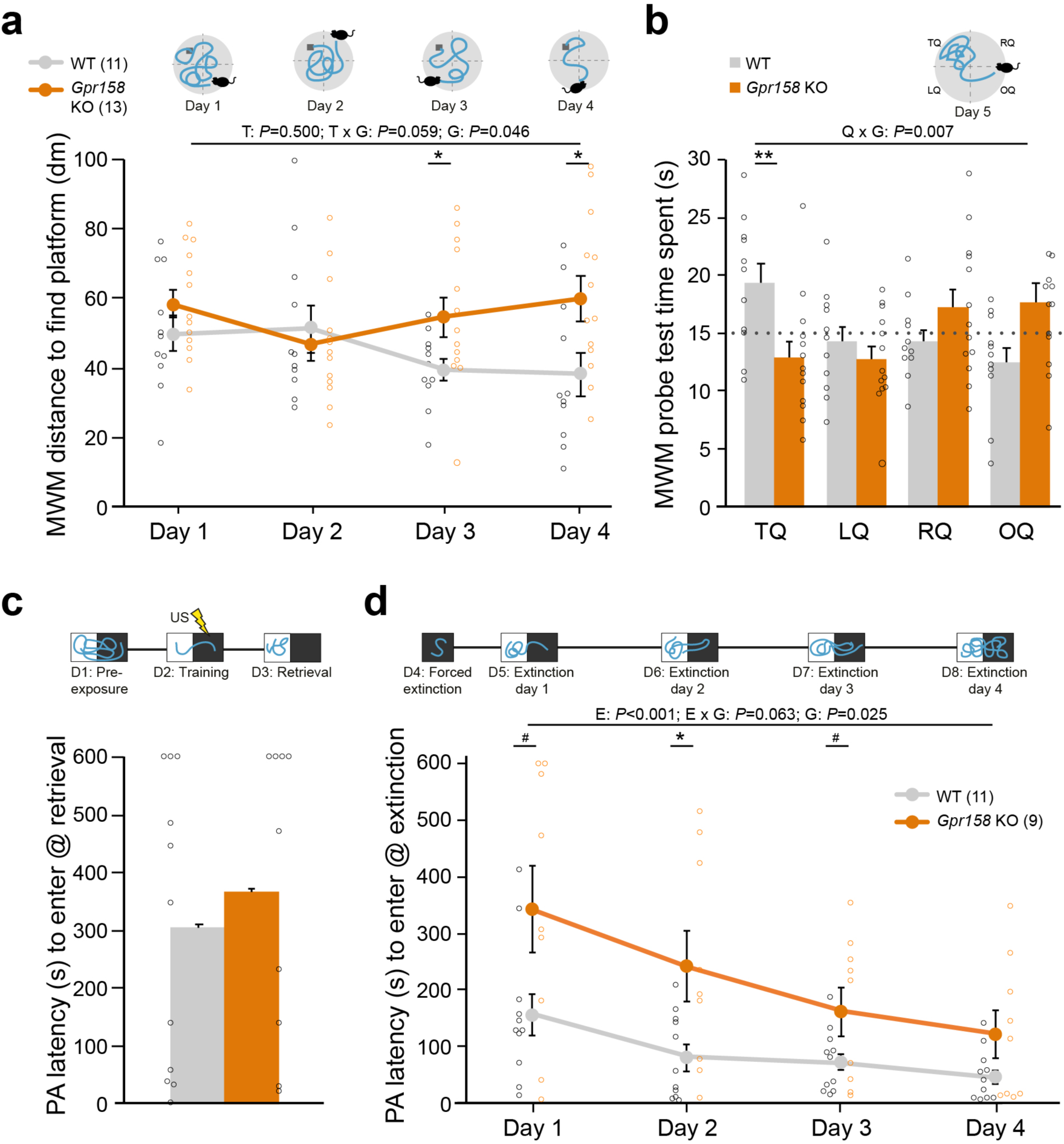
*Gpr158* KO have a spatial memory deficit and show delayed safety learning. **a,b**) Experimental set-up and data of training (see Figure 1 supplement 1; a, during four days of training, mice were placed in different locations in the circular pool and distance to reach the hidden platform (dark grey) was recorded), and probe test (b, time spent in each quadrant was recorded) in the Morris water maze (MWM). **a**) During MWM training, *Gpr158* KO mice swam a longer distance to reach the platform and their performance did not reach WT levels (nWT=11, nKO=13; genotype (G) effect *P*<0.046; training (T) effect *P*=0.500, interaction *P*=0.059). **b**) *Gpr158* KO mice showed a long-term memory deficit during the probe test (quadrant (Q) x genotype (G) *P*=0.007), and spent less time in the target quadrant (TQ, RQ: right quadrant, LQ: left quadrant, OQ: opposite quadrant). **c,d**) Experimental set-up and data of PA test and latency to enter the dark compartment after training (c) and during the four days after forced extinction (d). **c**) *Gpr158* KO mice (n=9) showed similar PA retention as WT mice (n=11). **d**) After forced extinction in the dark compartment, *Gpr158* KO mice showed a delay in extinction learning (Extinction (E) *P*<0.001, genotype (G) *P*=0.025, interaction *P*=0.063). Data are presented as mean±SEM with individual data points indicated. Asterisks and octothorpe indicate the level of significance between WT and KO assessed by Student’s t-test/MWU test (Figure 1 supplement 2), ^#^ *P*≤0.100; * *P*≤0.050; ** *P*≤0.01.

Long-term memory was assessed with a 24 h retention interval after training during a probe session in which the platform was removed (Figure 1 supplement 1). As the probe test measures the preferred location in a fixed time window it is able to dissociate memory from general activity, and as such it is relatively insensitive to altered swimming speed (Vorhees and Williams, 2006). *Gpr158* KO mice spent significantly less time in the target quadrant (TQ), indicative of long-term spatial memory impairments (**Figure 1b**, mixed ANOVA: interaction quadrant x genotype *P*=0.007; t-test: TQ *P*=0.008).

To distinguish between disturbed spatial processing deficits and motor or visual impairments possibly contributing to the observed changes, a visual platform test was performed 1 month after the probe test using a cue to signal the platform. Latency to find the platform was the same for *Gpr158* KO and WT mice during the visual platform trials (WT=7.1±1.1 s, *Gpr158* KO=10.2±2.0 s; t-test, *P*=0.201, data not shown). Additionally, when tested in the open field arena, anxiety-like behavior measured as distance moved and anxiolytic-like behavior measured as time spent in the center, was not different between genotypes (Figure 1 supplement 1). Taken together, in the absence of motor and visual deficits or anxiety-like behavior, *Gpr158* KO mice exhibited spatial memory deficits in the MWM paradigm.

To assess context-specific memory, an independent batch of *Gpr158* KO and WT mice underwent a contextual fear conditioning (CFC) paradigm (Figure 1 supplement 1). *Gpr158* KO mice did not have any fear memory deficit, exhibiting equal distance moved and freezing as WT during the retrieval test 24 h after conditioning (Figure 1 supplement 1); distance: t-test *P*=0.715; freezing: t-test *P*=0.693). To corroborate these results and to assess extinction of the acquired aversive memory we probed passive avoidance (PA) behavior, in an independent batch of mice, in which mice learn to avoid the preferred dark environment where a foot shock was delivered (**Figure 1c,d**). Long-term avoidance memory was tested on day 3, after pre-exposure (day 1) and training (day 2, **Figure 1c**). Both *Gpr158* KO and WT mice showed the same level of retention (MWU *P*=0.656), corroborating the lack of contextual aversive memory deficits in *Gpr158* KO mice. Additionally, we assessed whether acquisition of extinction memory was different between genotypes (**Figure 1d**). To this end, we first performed a forced extinction session in the dark compartment in order to promote safety learning over response extinction only (Micale et al., 2017), and thereafter measured the latency to enter the dark compartment during the subsequent four days of extinction. This revealed a significant delay in extinction memory acquisition for *Gpr158* KO mice (**Figure 1d**, mixed ANOVA: Extinction *P*<0.001, genotype *P*=0.025, interaction *P*=0.063; t-test: day 1 *P*=0.054, day 2 *P*=0.020; day 3 *P*=0.077). Taken together, whereas *Gpr158* KO mice show no deficits in aversive memory acquisition, they do exhibit spatial memory deficits and difficulty in acquiring extinction memory, in the absence of motor and visual impairments or anxiety.

### *Normal basal excitatory and inhibitory synaptic transmission in* Gpr158 *KO mice*

Abnormal synaptic transmission can alter information processing, affect learning and memory, and impact behavior (Mayford et al., 2012; Mitsushima et al., 2013). To that end, spontaneous excitatory and inhibitory post-synaptic currents (sEPSC and sIPSC) were recorded from CA1 pyramidal neurons from WT and *Gpr158* KO mice after completion of behavioral tasks (**Figure 2a-d**, Figure 1 supplement 1). Basal sEPSC frequency (t-test *P*=0.193) and amplitude (MWU *P*=0.403) was comparable between genotypes (**Figure 2a,b**). Although the decay time of sEPSCs was found prolonged in *Gpr158* KO cells (t-test *P*=0.047; Figure 2 supplement 1), further analysis of the weighted tau of decay did not reveal any difference between genotypes (WT 9.79±1.42 ms, KO 10.56±1.91 ms, t-test *P*=0.294, data not shown). Finally, rise time of sEPSCs was equal for both groups (t-test *P*=0.179). Additionally, we did not observe any differences in the frequency (t-test *P*=0.470) of sIPSCs (**Figure 2c,d**), however a trend for a reduction in amplitudes was identified (MWU *P*=0.058). Inhibitory receptor kinetics were equal between genotypes (t-test, decay *P*=0.995, rise *P*=0.314). Taken together, our data from CA1 pyramidal neurons do not reveal any substantial difference in excitatory and inhibitory synaptic neurotransmission that could alter neuronal communication and impact behavior.

**Figure 2.**
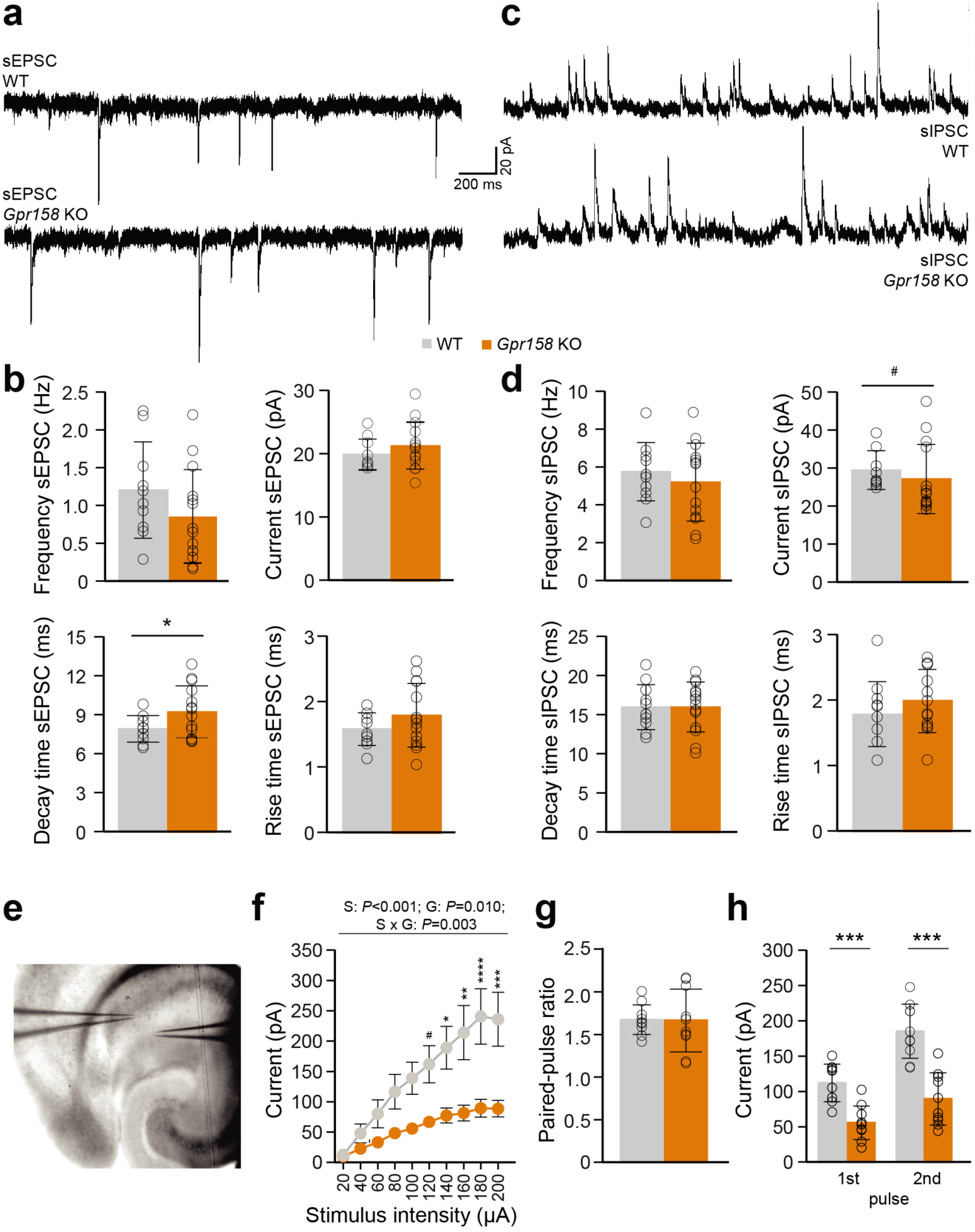
Electrophysiological assessment of *Gpr158* KO CA1 pyramidal incoming synaptic transmission. **a-d**) A cesium-gluconate-based intracellular was used to record both excitatory (a,b, WT n=10 cells from 4 animals, KO n=14 cells from 4 animals) and inhibitory (c,d, WT n=11 cells from 4 animals, KO n=14 cells from 4 animals) postsynaptic currents from CA1 pyramidal cells. Example traces of excitatory (a) and inhibitory activity (c) recorded from WT and *Gpr158* KO mice. Frequency, amplitude, and rise time of sEPSCs were equal between genotypes (b). A small increase in sEPSC decay time was observed in *Gpr158* KO (see Figure 2 supplement 1). No difference was observed in sIPSC frequency, decay time, or rise time (d). However, a trend for decreased sIPSC amplitude was observed in *Gpr158* KO (see Figure 2 supplement 1). **e-h**) For PPR recordings, a potassium-gluconate based intracellular was used for the recording pipette, while the stimulating pipette was loaded with aCSF and placed in a location where a clear unisynaptic response could be generated in the postsynaptic cell. Example configuration for I/O curve and PPR recordings (e; WT n=10 cells from 4 animals, KO n=10 cells from 3 animals). An I/O stimulation curve was generated by gradually increasing stimulation intensity in steps of 20 μA (f). A clear reduction in postsynaptic responses was observed in *Gpr158* KO upon Schaffer collateral stimulation (stimulus (S): *P*<0.001, genotype (G) *P*=0.010; interaction *P*=0.003) that became significant from 140 μA onwards. A succession of two stimulations separated by 50 ms was applied at stimulation intensity approximately half-maximal to that generated during the I/O curve. The ratio between the 1^st^ and 2^nd^ response amplitude was equal between genotypes (g). The raw amplitudes recorded during half-maximal stimulation during the PPR protocol differed significantly (h). For panel (f) data are presented as mean±SEM. For all other panels data are presented as mean±SD with individual data points indicated. Asterisks and octothorpe indicate the level of significance between WT and KO assessed by Student’s t-test or MWU (Figure 2 supplement 1), ^#^ *P*≤0.100; * *P*≤0.050; *** *P*≤0.001.

### *Normal release probability but reduced Schaffer collateral-mediated responses in* Gpr158 *KO mice*

A central feature in synaptic transmission is its ability to adapt to rapidly successive stimuli, which is regulated mainly by presynaptic mechanisms (Zucker and Regehr, 2002). The frequency of stimulation combined with synapse-specific release probabilities can filter activity, by allowing information transfer only when the appropriate conditions are met (Abbott et al., 1997; Fortune and Rose, 2001; Zucker and Regehr, 2002). To determine the excitability of CA3 to CA1 connections, an input-output (I/O) stimulation curve was generated, by stimulating Schaffer collaterals (SC) with increasing intensity, while recording from CA1 pyramidals (**Figure 2e**). *Gpr158* KO responses exhibited robust reduction for nearly all stimulation intensities, reaching plateau at below 100 pA, while WT responses plateau at just below 250 pA (**Figure 2c**, 2W-RMANOVA: genotype *P*=0.01, stimulation *P*<0.001, interaction *P*=0.003; Figure 2 supplement 1). The reduction in responses was not due to differences in the placement of the stimulating electrode, since the distance between recording and stimulating electrodes was equal between groups (WT 77.51±19.13 px, KO 86.39±18.46 px, *P*=0.305, data not shown). Subsequently, two successive stimuli were delivered, separated by 50 ms, at half-maximal stimulation intensities; current injected was not significantly different between genotypes (WT 90±21.60 µA, KO 102±12.76 µA, *P*=0.247, data not shown). Surprisingly, the paired-pulse ratio (PPR), was equal between the two genotypes (**Figure 2g**, t-test *P*=0.945), however the raw amplitudes recorded during the PPR protocol where robustly different between WT and *Gpr158* KO mice (**Figure 2h**, t-test, 1^st^ Pulse *P*<0.0001, 2^nd^ Pulse *P*<0.0001). Therefore, our data suggest that even though presynaptic release probability and machinery could be intact, compromised SC integrity, synapse numbers, or vesicle load could drive the attenuated SC-CA1 responses.

### Gpr158 *is implicated in hippocampal neuronal morphology*

Dendritic architecture is central to the neuron’s capacity to integrate signals, can influence cell intrinsic properties, and thus can impact learning and memory (Bekkers and Häusser, 2007). To that end, we investigated neuronal morphology both *in vitro* and *ex* vivo, by analyzing hippocampal primary culture morphology after *Gpr158* knock down (KD) (**Figure 3**), and by analyzing biocytin-loaded CA1 pyramidals in *Gpr158* KO and WT hippocampal slices (**Figure 4**).

**Figure 3.**
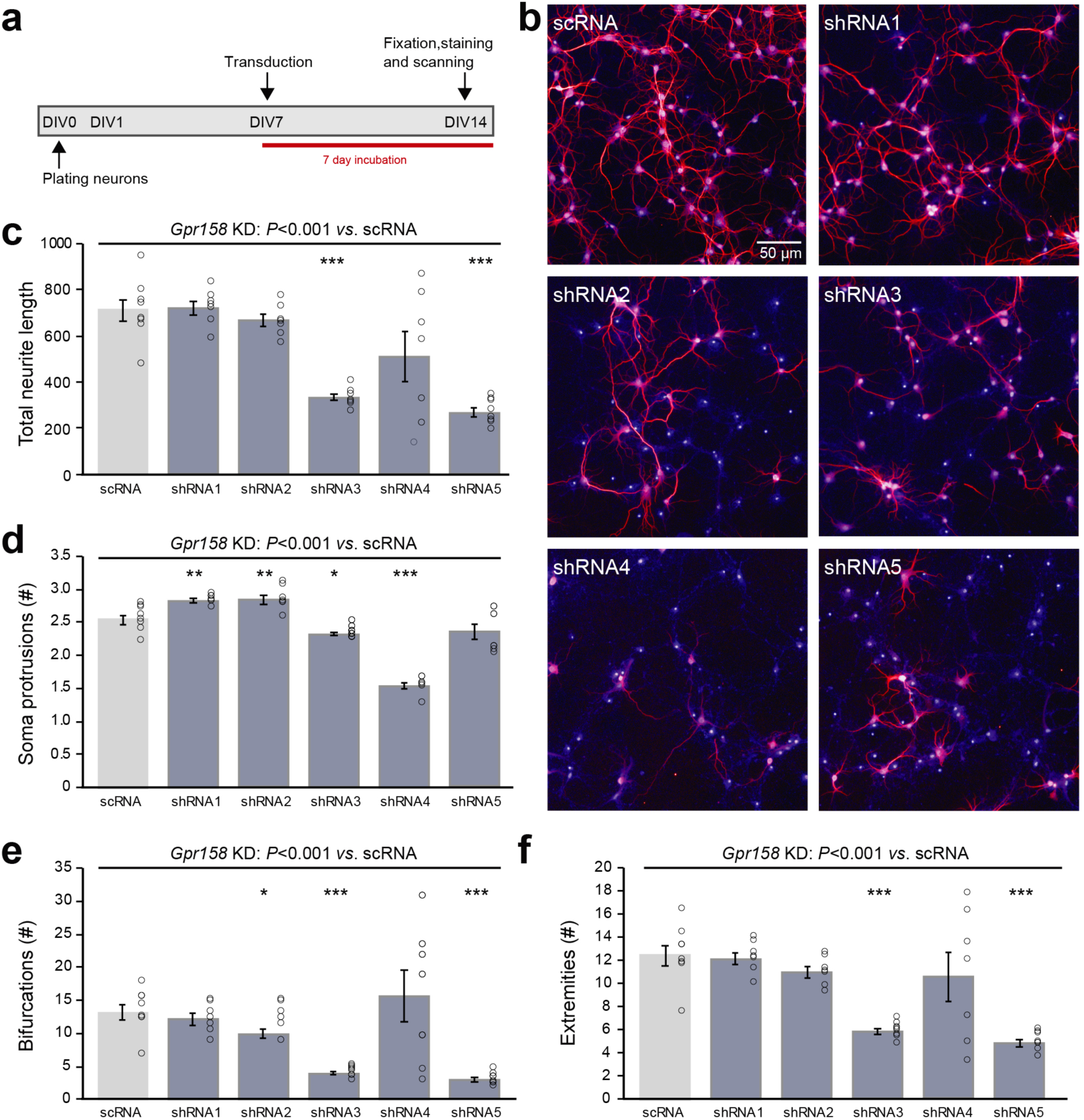
*Gpr158* knock down leads to reduced neuronal outgrowth of hippocampal primary neurons. **a**) Experimental set-up of *in vitro Gpr158* knockdown. Viral transduction with independent shRNAs occurred from DIV7 to DIV14, and morphological analysis occurred at DIV14. **b**) Representative images for control (scRNA) and *Gpr158* KD (shRNA1–5) neurons (red: MAP2, blue: nuclei). **c–f**) Morphological analysis after *Gpr158* knockdown (KD) at DIV14 (n=7–8 wells; see Figure 3 supplement 1) shows an overall effect of the shRNA treatment for all parameters (neurite length (c), number of soma protrusions (d), number of bifurcations (e) and number of extremities (f)), as tested by Kruskal-Wallis (*P*<0.001). Whereas shRNA3 affected all 4 parameters, the other shRNAs had a more limited effect for 1–3 of these parameters. Apart from a significant downregulation in number of protrusions, shRNA5 gave a similar strong effect as shRNA3, indicative of a *Gpr158*-related effect. Data are presented as mean±SEM with individual data points indicated. Asterisks indicate significant differences compared with control (scRNA) assessed by Student’s t-test (Figure 3 supplement 2), * *P*≤0.050; ** *P*≤0.010; *** *P*≤0.001.

**Figure 4.**
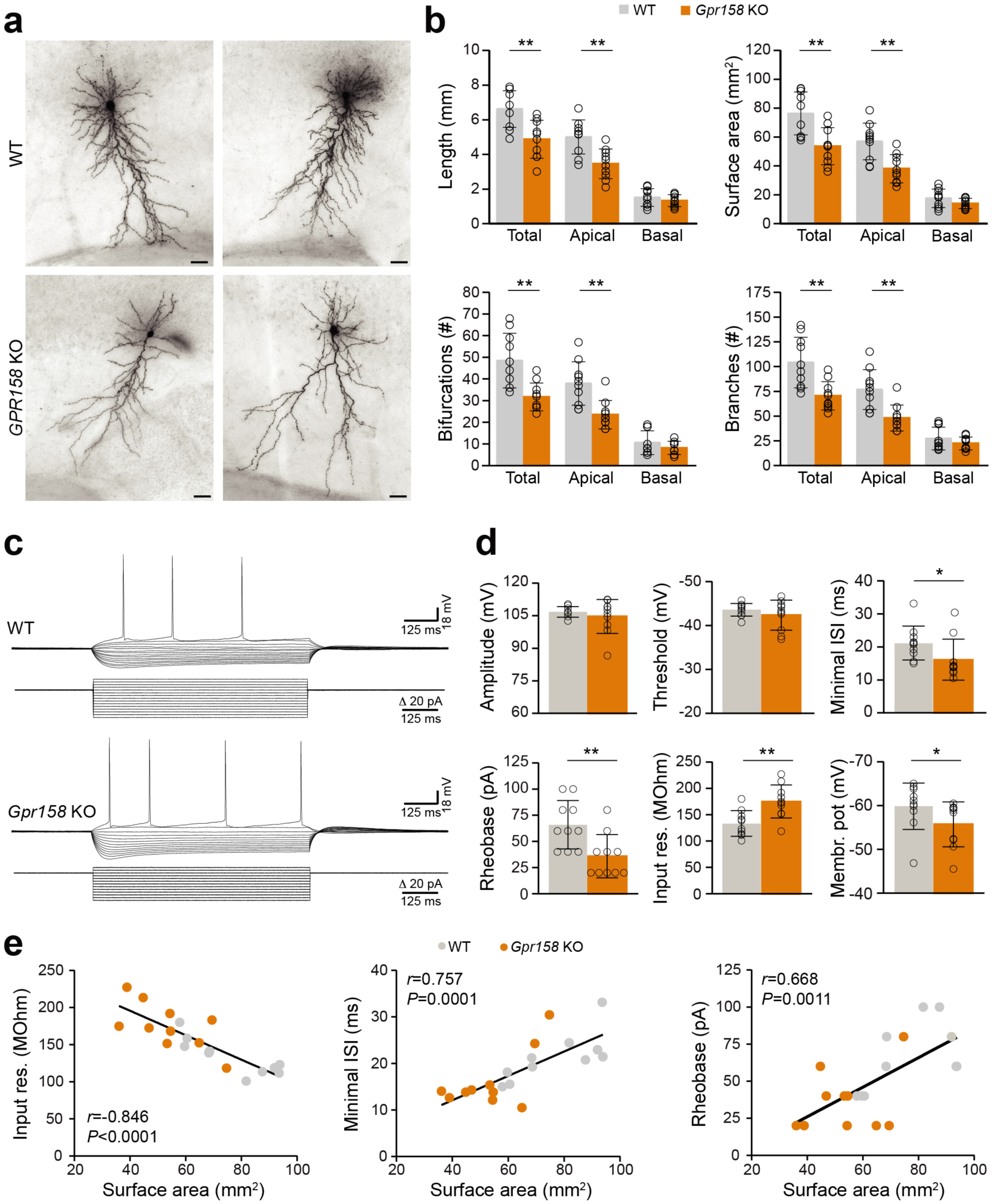
Reduced dendritic architecture supports hyperexcitability of *Gpr158* KO CA1 pyramidal neurons. **a**) Examples of CA1 pyramidal cells loaded with biocytin during electrophysiological recordings from WT (n=10 cells from 4 animals) and *Gpr158* KO (n=10 cells from 3 animals) hippocampal slices. Cells were subsequently reconstructed and their basic morphological measures were extracted and analyzed (Figure 4 supplement 1). **b**) Total dendritic length and surface area were significantly reduced in *Gpr158* KO CA1 pyramidals and the effect was specific to the apical compartment. A significant reduction in total and apical but not basal compartments was documented for the number of bifurcations and number of branches. **c–d**) For each reconstructed cell, the full action potential profile was generated and analyzed. **c**) Example traces of WT and *Gpr158* KO action potential profiles generated by current injections in steps of 20 pA. **d**) Action potential amplitude and threshold was unaffected by genotype. Minimum inter spike interval and rheobase were significantly reduced in *Gpr158* KO pyramidal cells. In addition, input resistance was increased in *Gpr158* KO and the membrane potential was more depolarised in *Gpr158* KO pyramidal cells. **e**) Significant Pearson correlations (*r*) and linear regression (Figure 4 supplement 2) between cell surface area and input resistance (*r*=-0.846, *P*<0.0001), minimum inter spike interval (*r*=0.757, *P*=0.0001), and rheobase (*r*=0.668, *P*=0.0011). Data are presented as mean±SD; individual data points are indicated. Asterisks indicate significant differences compared with WT cells assessed by Student’s t-test or MWU (Figure 4 supplement 1; Figure 4 supplement 2), * *P*≤0.050; ** *P*≤0.010.

Hippocampal primary neurons were transduced from day *in vitro* (DIV) 7 until DIV14, with one of five different shRNA constructs (shRNA1-5) targeting different parts of *Gpr158* mRNA sequence, or a control shRNA (scRNA) (**Figure 3a,b**). The window for transduction (DIV7–DIV14) was chosen based on the *in vitro* developmental gene expression of *Gpr158* (Figure 3 supplement 1), and the efficiency of shRNA-mediated KD of *Gpr158* by DIV14 (Figure 3 supplement 1). Four different parameters were measured underlying *in vitro* neuronal morphological development: total neurite length, number of protrusions from the soma, their bifurcations, and the number of extremities resulting from this (Figure 3 supplement 1). Although *Gpr158* KD reduced the survival of neurons for all shRNAs, shRNA4,5 were the strongest affected, while for shRNA1-3 neuronal survival was compromised less, hence allowing for full morphological analyses (Figure 3 supplement 1, Kruskal-Wallis *P*<0.001). All morphological parameters measured were affected by KD of *Gpr158* (**Figure 3c-f**, Kruskal-Wallis, all *P*<0.001; Figure 3 supplement 2). Specifically, neurite length (**Figure 3c**) was reduced by shRNA3 (t-test *P*<0.001) and shRNA5 (t-test *P*<0.001). KD of *Gpr158* resulted in variable effect on number of protrusions from the soma, with shRNA1 and shRNA2 yielding more protrusions (**Figure 3d**, t-test: shRNA1 *P*=0.004, shRNA2 *P*=0.010), shRNA3 and shRNA4 yielding a reduction in protrusions (**Figure 3d**, t-test: shRNA3 *P*=0.013, shRNA4 *P*<0.001), and shRNA5 having no significant effect (MWU: *P*=0.234). The number of bifurcations (**Figure 3e**) was reduced by shRNA2 (t-test, *P*=0.039), shRNA3 (t-test, *P*<0.001), and shRNA5 (t-test, *P*<0.001). Finally, the resulting number of extremities (**Figure 3f**) was reduced by shRNA3 (t-test, *P*<0.001) and shRNA5 (t-test, *P*<0.001). Taken together, in the reduced conditions of primary culture, *Gpr158* KD disturbed development of neuronal morphology.

### Polarized reductions in Gpr158 KO CA1 pyramidal cell dendritic architecture

Given the impact that *Gpr158* KD had on neuronal morphology *in vitro*, we subsequently reconstructed the dendritic morphology from biocytin-loaded CA1 pyramidal neurons in hippocampal slices from adult *Gpr158* KO and WT mice (**Figure 4a**). In line with the observations *in vitro, Gpr158* KO CA1 pyramidals exhibited reduced total dendritic length (**Figure 4b**, t-test *P*=0.002), as well as total surface area (**Figure 4b**, t-test *P*=0.002; Figure 4 supplement 1). Moreover, the observed reductions in total length and surface area were supported by reductions specifically in the apical but not the basal dendritic compartment (**Figure 4b**, t-test: length [apical *P*=0.001, basal *P*=0.346], surface [apical *P*=0.002, basal *P* =0.136]). Furthermore, the general complexity of dendritic arborizations in *Gpr158* KO was reduced, reflected by the decrease in the total number of bifurcations per cell (**Figure 4b**, t-test: total *P*=0.002, apical *P*=0.001, MWU: basal *P*=0.481). In contrast to the variable effect observed *in vitro*, the number of dendrites emerging from the cell body was equal (WT 7.40±1.78, KO 7.70±1.95, t-test *P*=0.723, data not shown). The total number of branches was significantly reduced in *Gpr158* KO and the effect was specific to the apical dendritic compartment (**Figure 4b**, t-tests: total *P*=0.003, apical *P*=0.001; MWU: basal *P*=0.529). Thus, the biocytin reconstructions of *Gpr158* KO CA1 pyramidal neurons *ex vivo,* illustrated reductions in dendritic architecture, paralleling *in vitro* observations, and localized the effect specifically to the apical dendritic compartment.

### Reduced dendritic architecture associates with augmented intrinsic excitability of *Gpr158* KO CA1 pyramidal cells

Neuronal morphology contributes substantially to the integrative properties of neurons, and can have a major influence on neuronal firing by altering intrinsic cell properties (Bekkers and Häusser, 2007; Kowalski et al., 2016). Given the robust changes in neuronal architecture we documented both *in vitro* and *ex vivo*, we analyzed the action potential (AP) profiles from WT and *Gpr158* KO CA1 pyramidal cells in hippocampal slices (**Figure 4c**) used for morphological assessment. AP amplitude (**Figure 4d**, *P*=0.439, Figure 4 supplement 2) and threshold (**Figure 4d**, *P*=0.311) did not differ for genotype. However, both the minimum inter spike interval (**Figure 4d**, ISI, *P*=0.015) and minimum current required to elicit the first AP (**Figure 4d**, rheobase, *P*=0.009) were reduced, underlying increased excitability of *Gpr158* KO pyramidals. Moreover, the input resistance (Ri) of *Gpr158* KO cells was significantly increased, further substantiating and promoting their increased excitability (**Figure 4d**, *P*=0.004). Neuronal cell membrane tau did not differ between the two groups (WT: 29.92±4.84 ms, KO: 27.77±4.42 ms, *P*=0.312, data not shown). Finally, the resting membrane potential of *Gpr158* KOs was found more depolarized than in WT (**Figure 4d**, *P*=0.035).

We then applied principal component analysis (PCA) on the morphological and electrophysiological parameters collected from the same cells to dissect the most significant determinant underlying the changes observed (Figure 4 supplement 2). For all components, cell surface area was the variable with the most significant contribution (Figure 4 supplement 2, first 3 components shown). A subsequent correlation matrix of surface area against all other morphological and electrophysiological variables demonstrated significant correlations with all morphological measures, with cell excitability, and with PPR raw amplitudes (Figure 4 supplement 2). However, upon multiple comparison correction, the correlation between surface area and raw PPR amplitudes did not retain significance, while the rest retained high statistical significance (Figure 4 supplement 2). Subsequent linear regression demonstrated that surface area negatively correlated with Ri (**Figure 4e**, *r*=-0.846, *P*<0.0001), whereas a positive relationship was demonstrated with minimum ISI (**Figure 4e**, *r*=0.757, *P*<0.001), and rheobase (**Figure 4e**, *r*=0.668, *P*=0.001). Taken together, our data demonstrated that *Gpr158* KO CA1 pyramidal cells exhibit increased excitability, and that this increase is likely driven by their notable reduction in dendritic architecture.

### Reduced architecture of CA1 pyramidal neurons supports MWM learning deficits

Changes in the excitability of neurons can alter information flow, whereas changes in dendritic architecture can also impact the integrative capacity of neurons (Bekkers and Häusser, 2007; Magee, 2000; van Elburg and van Ooyen, 2010). Moreover, dendritic complexity appears to increase with evolution and with increasing cognitive demands, with human prefrontal pyramidals exhibiting the largest cortical dendritic complexity (Elston et al., 2001; Mohan et al., 2015). Furthermore, increasing dendritic complexity of human cortical pyramidals appears correlated with increasing intelligence quotient, in a small sample of surgically resected brain tissue (Goriounova et al., 2018). Since the electrophysiological datasets reported here were generated from mice that underwent behavioral assessment (Figure 1 supplement 1), we then averaged the morphology data of all cells collected to generate an overview of dendritic architectures for n=6 animals per genotype (**Figure 5a**). Analysis of the combined morphologies further substantiated the robust reduction in dendritic architecture of *Gpr158* KO hippocampal neurons (**Figure 5a**, 2W-ANOVA, genotype (G) effect; length *P*=0.002; surface *P*<0.001; bifurcations *P*<0.001; branches *P*=0.001; Figure 5 supplement 1). Furthermore, the effect was specific to the apical but not the basal dendritic compartment (**Figure 5a**, 2W-ANOVA, dendritic compartment (DC) effect; for all measures *P*<0.001). Apical (*P*<0.001) but not basal dendritic length (all *P*>0.300) was reduced in *Gpr158* KO mice, as well as surface area, number of branches, and number of dendritic bifurcations in the same manner (Figure 5 supplement 1).

**Figure 5:**
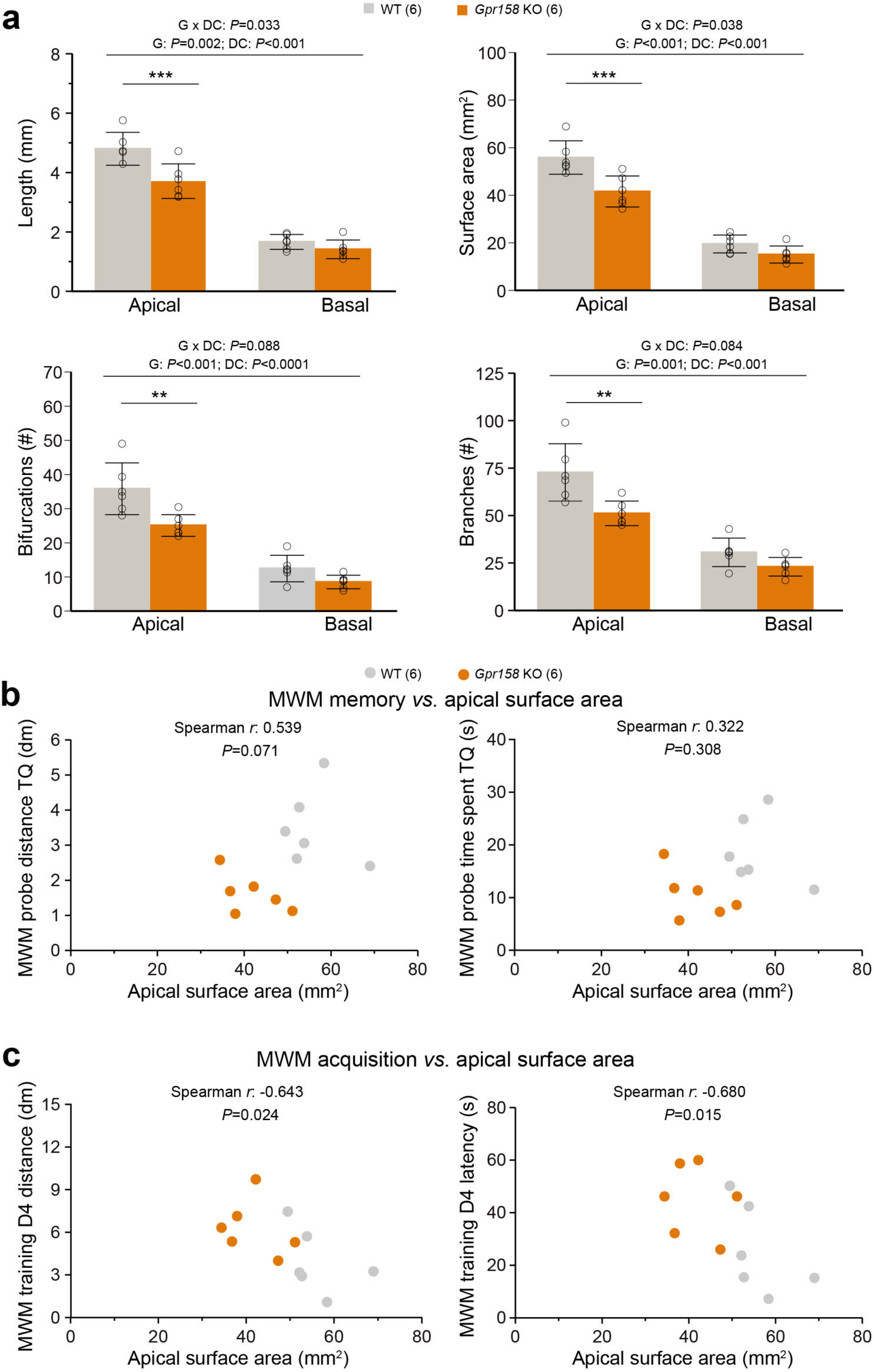
Reduced dendritic architecture negatively impacts learning during MWM training. **a**) Cell morphology from all the electrophysiological data sets described were averaged to generate an overview of hippocampal dendritic architecture per animal (WT n=6 animals, 18 cells averaged, KO n=6 animals, 17 cells averaged). Apical but not basal hippocampal dendritic length, surface area, bifurcations and branches were significantly reduced in *Gpr158* KO animals. Apical surface area did not (significantly) correlate with either the distance covered (*P*=0.071) or the time spent (*P*=0.308) in the target quadrant (TQ) during the MWM probe test assessing long-term memory. **c**) During acquisition of the MWM task, the apical surface area did significantly and negatively correlate with both the distance covered to reach the platform (*P*=0.024) and the latency (*P*=0.015) to reach the platform on the last training day (D4). Data are presented as mean±SD with individual data points indicated. Asterisks indicate significant differences between WT and KO assessed by Student’s t-test (Figure 5 supplement 1), ** *P*≤0.010; *** *P*≤0.001.

Given the emergence of surface area as major determinant for the morphological and electrophysiological differences observed, we further probed its relationship with MWM parameters for the entire group of mice. For the MWM probe test, neither the distance covered in the TQ (**Figure 5b**, Spearman *r*=0.538, *P*=0.071), nor the time spent in the TQ (**Figure 5b**, Spearman *r*=0.322, *P*=0.308) were significantly correlated with the animal’s apical dendritic surface area. However, for the acquisition of the MWM task, surface area did significantly and negatively correlate with both the distance covered to reach the platform (**Figure 5c**, Spearman *r*=-0.643, *P*=0.024), and with the latency to reach the platform (**Figure 5c**, Spearman *r*=-0.680, *P*=0.015). Therefore, the reduction in general dendritic architecture in *Gpr158* KO CA1 neurons, although not directly affecting long-term MWM memory, imposed a significant and negative correlation with the acquisition of this spatial memory task.

## Discussion

In this study we report that mice deficient of *Gpr158* exhibit impairments in the acquisition and retrieval of spatial memory in the MWM paradigm, as well as deficits in the learning of an extinction memory in the PA task. Although hippocampal CA1 basal excitatory and inhibitory neurotransmission was found normal, we report a remarkable reduction in the excitability of SC synapses onto CA1 pyramidals, whereas we observed a potential compensatory increase in the intrinsic excitability of these neurons. Strikingly, both *in vitro* and *ex vivo* we observe a large effect of reduced *Gpr158* levels on dendritic morphology. *Ex vivo*, the dendritic architecture reductions of CA1 neurons was specific for the apical but not the basal compartment of these cells. Finally, the reduced apical complexity of these neurons appears to support the spatial memory acquisition deficits in these mice, since reduced neuronal surface area positively correlated with poor performance during acquisition of the MWM task.

### *Acquisition of a safety memory and* s*patial memory are affected in* Gpr158 *KO mice*

Non-spatial memory acquisition and expression, as measured in the passive avoidance test, and in contextual fear conditioning with context as the foreground stimulus, was not affected in *Gpr158* KO mice. However, normalization of avoidance behavior after forced exposure to the conditioning context was delayed in *Gpr158* KO mice. This safety learning procedure has been shown to rely on hippocampus CB1 receptors (Micale et al., 2017), in contrast to response extinction learning, i.e. repeated execution of the avoided response in absence of negative consequences, that is largely cortically mediated. Single intrahippocampal injection of cannabinoid-related compounds prior to safety learning acquisition could alter the effect of safety learning (Micale et al., 2017). Additionally, these compounds affected the propagation of signaling from the DG via area CA3 to the CA1 subfield upon perforant pathway stimulation (Micale et al., 2017). Thus, the reduced SC-mediated responses we observed in hippocampal slices of *Gpr158* KO mice might indicate that information processing within the trisynaptic circuitry of the hippocampus is disturbed. As such this precipitates not only in impaired safety learning, but also in poor spatial memory acquisition and expression, which is also dependent on balanced cannabinoid signaling in the hippocampus (Abush and Akirav, 2010).

During training in the MWM paradigm, *Gpr158* KO mice exhibited robust deficits in task acquisition, demonstrated by their increased distance covered and time spent to reach the platform. This is in line with a previous observation of impaired MWM acquisition in these mice (Khrimian et al., 2017). We additionally report that *Gpr158* KO mice demonstrated a long-term spatial memory deficit, highlighted by the reduced performance during the MWM probe trial. During the probe test, the platform is removed and the animal is requested to rely on spatial information to relocate the area of the previous escape platform after a 24 h interval. The hippocampus plays a major role in spatial learning, with involvement of both the CA1 and CA3 regions (Florian and Roullet, 2004), in which *Gpr158* is expressed (Condomitti et al., 2018; Khrimian et al., 2017). Theta frequency coupling between CA3 and CA1 is enhanced during MWM learning, when there is a clear improvement in task performance (Hernández-Pérez et al., 2016). The changes in CA1 morphology and excitability we report along with the changes in CA3 physiology (Condomitti et al., 2018) could alter such coupling, hence impacting learning in *Gpr158* KO mice. Although the CA3 region is involved in spatial memory acquisition, it is not required for long-term memory recall during the probe test (Florian and Roullet, 2004) that we here report as also being affected in *Gpr158* KO mice. Enhanced CA1 LTP, is shown to improve MWM probe test performance (Okada et al., 2003). Although not yet demonstrated, LTP deficits in the CA1 region akin to those described in CA3 (Condomitti et al., 2018) are likely - as discussed below, and could support deficits in spatial memory acquisition.

### Reduced dendritic architecture supports spatial memory learning deficits

Central to our observations are the striking reductions in dendritic architecture and complexity of *Gpr158* deficient neurons, both *in vitro* - in hippocampal cultures, and *ex vivo -* in reconstructed CA1 pyramidals within hippocampal slices. Complex dendritic architecture is essential to proper learning and memory (Elston et al., 2001). Dendritic complexity increases with evolution and with increasing cognitive demands (Tronel et al., 2010), and impairments in dendritic architecture have been observed in disorders with reduced cognitive capacities (Kulkarni and Firestein, 2012). Moreover, dendritic structure impacts the integrative ability of neurons, through pathway-specific distinct input domains, coincidence detection, synaptic scaling of distal inputs, and targeted dendritic inhibition, to name a few (Spruston, 2008).

Importantly, the reduced architecture we observed, negatively correlated with the last days of spatial memory acquisition in in the MWM paradigm but not significantly with probe test performance. Spatial learning itself has been shown to enhance dendritic complexity by increasing length, bifurcations, and branches of DG new born granule cells, in rats trained in MWM and in the delayed matching-to-place task (Tronel et al., 2010). Mostly these learning-induced changes are relatively small and short-lasting. Instead, genetic-induced mutations that affect dendritic architecture during development could have a higher impact. Hence mutation-associated learning deficits could arise through changing specific plasticity mechanisms. Mutant mice for tyrosine kinase *Fyn*, involved in normal cell physiology, exhibit deficits in MWM acquisition and recall, along with impairments in CA1 LTP, and reduced compaction and organization of CA1 apical dendrites (Grant et al., 1992). In addition, mice double mutant for cysteine proteases calpain 1 and 2 exhibit impaired spatial memory in MWM, in the absence of visual or motor deficits or anxiety-like behavior in the open field (Amini et al., 2013). Electrophysiological analysis revealed that CA1 pyramidal dendritic complexity, LTP, and CA3 to CA1 I/O curve were robustly reduced in these mutants (Amini et al., 2013). It is therefore likely that the reduced dendritic architecture of *Gpr158* KO CA1 pyramidals we report here could faithfully underlie the deficits in spatial memory acquisition, potentially through impairments in LTP.

In the SC to CA1 network, brain-derived neurotrophic factor (BDNF) is required for the induction and maintenance of LTP brought about by repeated dopamine stimulation in apical but not in basal dendrites (Navakkode et al., 2012), and hippocampal BDNF levels in *Gpr158* KO are found reduced (Khrimian et al., 2017). To date, impaired PPR and LTP in *Gpr158* KOs has been reported for the MF to CA3 pathway (Condomitti et al., 2018; Khrimian et al., 2017), but LTP has not been investigated for the CA3 to CA1 pathway. Although we report normal PPR for this pathway, given the robust reduction in CA3 to CA1 stimulation and deficient dendritic architecture we observe, along with reduced hippocampal BDNF levels (Khrimian et al., 2017), it is safe to speculate that CA3 to CA1 LTP will likely be impaired in *Gpr158* KO as well, driving impairments in spatial memory.

### Normal basal synaptic transmission and increased excitability of *Gpr158* KO CA1 pyramidals

Along with the changes in dendritic architecture, we observed an overall increase in the excitability of the reconstructed cells. Rheobase as well as minimal ISI were significantly reduced in *Gpr158* KO CA1 pyramidals, whereas input resistance was increased. Moreover, PCA revealed that these changes were strongly correlated with the cell surface area. The contribution of the dendritic tree to neuronal excitability has been elegantly demonstrated, whereby pinching or severing neuronal dendrites significantly increased action potential frequency and input resistance, while reducing rheobase (Bekkers and Häusser, 2007). As such, our observations of increased CA1 pyramidal excitability in *Gpr158* KOs are indeed likely governed by their reduced dendritic morphology. This increased excitability could compensate for the reduced SC drive, as previously reported in an epilepsy model (Dinocourt et al., 2011). A hallmark of CA1 neurons is the ability to show action potential backpropagation, which is important for the integration of synaptic input and the induction of synaptic plasticity. Dendritic architecture, and specifically the number of dendritic branchpoints, showed a strong relationship with the functional threshold of backpropagation (Vetter et al., 2001), with lower number of branchpoints having a lower requirement of Na-channels to induce backpropagation as shown by computational modeling. As such, a lower threshold to induce backpropagation could compensate for the reduced SC drive and possible deficits in LTP.

Basal synaptic transmission onto CA1 pyramidal cells appears unaffected in *Gpr158* KO hippocampus, with no notable changes in either spontaneous excitatory or inhibitory frequency and amplitude observed, in line with recent observations for evoked IPSCs (Ostrovskaya et al., 2018). This is in contrast the reductions observed in sEPSC frequency and amplitude in *Gpr158* KO CA3 (Condomitti et al., 2018). Along with the reductions in evoked AMPA and NMDA CA3 currents (Condomitti et al., 2018) as well as the reduced SC mediated responses we observe, it is conceivable that when the network is further challenged, or when miniature release is assessed, such changes may become apparent in CA1 as well. Albeit, reduced CA3 to CA1 input does not always impact basal synaptic transmission (Amini et al., 2013; Grant et al., 1992).

### Is Gpr158 an inducer of presynaptic organization in CA1 akin to its role in CA3?

A major contributing factor in affecting *Gpr158* KO CA1 pyramidal morphology is incoming presynaptic activity. In the rodent hippocampus, mechanical perforant pathway denervation leads to sustained reductions in dendritic length and complexity of the postsynaptic dentate gyrus granule cells (Vuksic et al., 2011). Importantly, these reductions were observed primarily in the outer molecular layer where denervation had occurred. Additionally, deafness or facial nerve lesions resulted in pathway-specific reductions in dendritic morphology, in either primary auditory cortex (Bose et al., 2010), or primary motor cortex pyramidal cells (Urrego et al., 2015), respectively. The attenuated responses of SC stimulation we report, along with the reported reduction in CA3 pyramidal excitability (Condomitti et al., 2018), could lead to reduced SC output. Our reconstruction of *Gpr158* KO CA1 pyramidals revealed that structural reductions were specific to the apical but not basal dendritic compartments. Consequently, attenuated excitation of CA1 apical dendrites by SC might promote pathway-specific morphological alternations we report here. However, how this pathway-specificity is obtained is not clear, as CA3 SC project both to apical and basal dendrites. Albeit, Gpr158 could be required for the formation of SC to apical but not basal CA1 dendritic synapses.

In the MF-CA3 pathway, Gpr158 functions as a post-synaptic organizer (Condomitti et al., 2018), where it has a specific role in shaping synapse morphology and function in apical dendrites within the MF-CA3 pathway, but not within the CA3-CA3 recurrent pathway. This is largely ascribed to the enriched subcellular localization of Gpr158 to apical dendrites in the stratum lucidum (Condomitti et al., 2018). With the specific effect in the apical part of CA1 dendrites in *Gpr158* KO mice, one could assume that subcellular localization of Gpr158 in CA1 neurons is similar to that of CA3 neurons. As such, it raises the possibility that the CA1 dendritic phenotype is independent of SC input and instead that SC input is induced by malformed synaptic contacts due to a lack of normally enriched levels of Gpr158. Our *in vitro* KD data also suggest a morphological role during development of Gpr158. Transduction of *Gpr158* shRNAs at hippocampal primary neurons at DIV7 showed at least in two out of five a major effect on dendritic outgrowth and maintenance in the weeks thereafter. Taken together, lack of the post-synaptic organizer *Gpr158* itself, and/or the resulting altered dendritic morphology, could subsequently shape presynaptic SC input.

Recently, to the growing set of post-synaptic organizers of the hippocampus CA1 (Thakar et al., 2017) another three important factors have been added recently (Schroeder et al., 2018), i.c. Flrt2, Lrrtm1, and Slitrk1. The authors elegantly showed that these factors affect synaptic organization in a different manner, and that they are expressed in overlapping sets of neurons, with a ∼23% probability of being expressed all together at the same synapse with PSD-95. *Gpr158* expression in CA1 has a temporal shift compared with that in CA3. Knowing that synapse formation starts at PD7 and continues until PD28, and that *Gpr158* expression in CA1 starts at PD10-14 and reaches adult-like levels in term of CA1/CA3 ratio at PD28 (Condomitti et al., 2018; Khrimian et al., 2017), Gpr158 could well function together with the growing set of CA1 postsynaptic organizers in a subset of SC-CA1 connections. An outstanding question remains which CA3 expressed presynaptic organizer interacts with postsynaptic Gpr158 to form CA3-CA1 synapses. From the recently reported 129 interactors (Orlandi et al., 2018), 12 were extracellular matrix-related, and of these several are expressed at low-high levels in CA3 (Supplementary file).

Taken together, we here showed that apart from the previously reported important role of Gpr158 in the MF-CA3 pathway, also the CA3-CA1 pathway is affected by loss of *Gpr158*. Specifically, we showed that CA1 dendritic morphology and synaptic function is related to the spatial learning deficit that *Gpr158* KO mice display.

## Animals, material and methods

### Animals

*Gpr158* knock-out (KO) mice (*Gpr158*^tm1(KOMP)Vlcg^) were generated by replacing the *Gpr158* gene with a LacZ cassette including a stop codon in the region of exon 1 and exon 2 (Orlandi et al., 2015), and were purchased from the KOMP repository (see Figure S1). This line was bred for >5 generations on a C57BL/6J background (Charles River, France) in the animal facility of the VU University Amsterdam in standard type-2 Macrolon cages, enriched with nesting material on a 12/12 h light/dark rhythm (lights on at 7:00 AM), a constant temperature of 23±1 °C, and a relative humidity of 50±10%. Food and water were provided ad libitum. After weaning, all mice were group-housed per sex. For behavioral experiments, male mice were single-housed 1-2 weeks prior to the first test, and received a PVC tube and wooden chew stick as additional enrichment.

For all behavioral experiments, we used male 8–12 weeks old *Gpr158* KO mice and wild type (WT) littermates as control. Electrophysiological recording and biocytin fillings were performed on the same males that were tested in the Morris water maze and open field, with an >8 week interval after the last test, at the age of ∼16-20 weeks (Figure 1 supplement 1). For primary hippocampal cultures, E18 embryos were taken from pregnant WT (C57BL/6J) mice. All experiments were performed in accordance to Dutch law and licensing agreements and protocols were approved by the Animal Ethics Committee of the VU University Amsterdam.

### Morris water maze (MWM) test

Animals were handled for one week, two times a day, prior to MWM training. A circular pool (125 cm in diameter) was filled with opaque-colored (nontoxic dye) water and the water temperature was kept at 23–25°C. A round transparent escape platform (9 cm in diameter) was placed in the top left quadrant (target quadrant) of the pool, hidden 0.2 cm below the water surface. Geometric visual cues (1 m from the pool) were located on the walls of a dimly lit room (20 lx).

Each mouse received a total of 2 daily swim sessions, each comprised of 2 swim trials (maximum duration 60 s per trial), with an interval of 2 minutes between sessions for 4 consecutive days. Before starting each training day, mice were placed on the platform for 30 s and then placed into the water at a semi-random start position (out of 4 start-positions). Mice that failed to find the platform were guided and placed on the platform for 15 s by the experimenter on day one. After each swim session, mice were returned to their home-cage for 2 minutes. Latency and distance swam to find the platform during training, and time spent and distance swam in the target quadrant in the probe test were examined using video tracking Viewer 2 (Biobserve, Bonn, Germany). Latency and distance to find the platform during training served as read-out of spatial memory acquisition. On the 5^th^ day, the probe test (60 s) was performed, in which the platform was removed and mice were placed into the water opposite to the location of the platform/target quadrant. The time spent in each quadrant was recorded and time spent in the target quadrant served as measure of long-term spatial memory.

### Passive avoidance test

The pre-exposure, training, retention test, forced-exposure and five-day extinction protocols were performed in a passive avoidance (PA) system (Model 256000, TSE-Systems, Bad Homburg, Germany). In the pre-exposure protocol, without unconditioned stimulus (US), mice were placed in the bright compartment (1000 lx) for 15 s. Then, a sliding door was opened for allowing the mice to explore the dark compartment (∼ 10 lx) for a maximum period of 300 s. The training and retention test were performed as described before (Baarendse et al., 2008) with minor modifications in the US intensity (0.7 mA) and duration (2 s). The forced-exposure protocol was performed depending on the maximum time spent in the dark compartment during training, totaling 600 s. The extinction protocol was performed as the pre-exposure protocol, but now for a maximum period of 600 s. During the retrieval and extinction protocols, the latency to enter the dark compartment was measured as a read-out of long-term memory. Mice that did not enter the dark compartment during retrieval and extinction tests were recorded as a latency of 600 s. Boxes were cleaned with 70% ethanol between each experiment.

### Contextual fear conditioning

Contextual fear conditioning was carried out in a fear conditioning system (TSE-Systems, Bad Homburg, Germany) as described before (Schmitz et al., 2017). Long-term memory was assessed 24 h after training.

### Open field

The open field test (OF) was performed in a white square box (50 × 50 cm, 35 cm high, 200 lx) by placing mice in the corner of the box and recording their exploration behavior for 10 minutes by video tracking (Viewer 2, Biobserve GmbH, Bonn, Germany). Time spent in the center area (32.9×32.9 cm) and total distance moved during the 10 minute test were used as parameters for anxiety-like behavior and locomotor performance. Open field boxes were cleaned with 70% ethanol between each experiment.

### Primary hippocampal culture, short hairpin RNAs and lentivirus production

Hippocampi were collected in Hanks balanced salts solution (HBSS; Sigma-Aldrich, St. Louis, MO) with 7 mM HEPES (Life Technologies/Gibco, Carlsbad, CA) and trypsin was added (10% final concentration; Life Technologies/Gibco) for 15-20 minutes at 37 °C. Following two washing steps with HBSS-HEPES and 1 time with neurobasal medium (supplemented with 2% B-27, 1.8% HEPES, 0.25% glutamax and 0.1 % penicillin/streptomycin (all from Life Technologies/Gibco). Cells were triturated with a fire-polished Pasteur pipette. After cell dissociation, neurons were plated at a density of 12.5 x 10^3^ cells/well in a 96-well plate (Cellstar, Greiner Bio-One, Frickenhausen, Germany) coated with poly-D-lysine and laminin (Sigma-Aldrich) and treated with 5% heat-inactivated horse serum (Life Technologies/Gibco), or at a density of 75 x 10^3^/well in a 12-well plate. Neurons were kept at 37 °C / 5% CO_2_.

Short hairpin RNAs (shRNAs) were purchased from Sigma-Aldrich (Mission^®^ shRNA bacterial glycerol stock; SHCLNG-XM_140850). Five shRNA were used to target and knockdown *Gpr158* gene expression and a scrambled shRNA (Sigma-Aldrich, pLKO.1-puro Non-Mammalian shRNA) was used as a control. Sequences for shRNA knockdown of the *Gpr158* gene were: shRNA#1 (TRC number: TRCN0000028697, sequence: CCGGGCCAAGTACATTTCGTTGTATCTCGAGATACAACGAAATGTACTTGGCTTTTT), shRNA#2 (TRC number: TRCN0000028716, sequence: CCGGGCTCATTATCACGGCTATATTCTCGAGAATATAGCCGTGATAATGAGCTTTTT), shRNA#3 (TRC number: TRCN0000028727, sequence: CCGGCCGGTCGTTATTCTGTACTTTCTCGAGAAAGTACAGAATAACGACCGGTTTTT), shRNA#4 (TRC number: TRCN0000028742, sequence: CCGGCGGCTATATTCCATACAATTACTCGAGTAATTGTATGGAATATAGCCGTTTTT) and shRNA#5 (TRC number: TRCN0000028751, sequence: CCGGCCTTAACAACTCAGAGTGTATCTCGAGATACACTCTGAGTTGTTAAGGTTTTT).

Plasmids of shRNAs were plated on Lysogeny broth (LB) agar medium supplemented with ampicillin (Sigma). Single colonies were picked and grown in LB liquid medium supplemented with ampicillin using a shaker incubator at 37 °C. DNA was extracted using NucleoBond Xtra Midi kit (Bioké, Leiden, The Netherlands) according to the manufacturer’s protocol. Lentivirus production were done using packaging, transducing and envelope constructs at the same time for human embryonic kidney 293T cells transfection. Lentiviral particles were collected after 2 days of transfection and concentrated using ultracentrifugation (Beckman Coulter, Optima^TM^ LE-80K).

Neurons were separately transduced with a *Gpr158*-targeting shRNA (see below) lentivirus at days in vitro 7 (DIV7). Briefly, neurons were transduced with 1:4000 titrated lentiviral particles in neurobasal medium supplemented with 2% B-27, 1.8% HEPES, 0.25% glutamax, 0.1% penicillin/streptomycin for each shRNA. Transduced neurons were kept at 37 °C / 5% CO_2._ Cells for immunostaining were fixed at DIV14 using 4% PFA, 2% saccharose (VWR chemicals) in PBS (pH 7.4) for 20 minutes. Cells for quantitative real time PCR (qRT-PCR) experiments were harvested using Trizol reagent (Life Technologies) at DIV14.

### Immunocytochemistry for neurite morphology analysis

Fixed neurons were washed with PBS and permeabilized with 0.5% Triton X-100 (Sigma Aldrich) in PBS (Life Technologies) for 10 minutes. After washing, cells were blocked with 1% BSA (Sigma Aldrich) and 0.1% Triton X-100 (Sigma Aldrich) in PBS for 1 h. Neurons were stained with chicken anti-MAP2 (Bio-Connect, Huissen, The Netherlands; 1:5,000) at 4 °C overnight, and visualized using anti-chicken Alexa Fluor 647 (Life Technologies/Gibco; 1:400) for 90 minutes at RT. The nucleus was stained with Hoechst dye (1:10,000 in distilled water) for 10 minutes at RT.

### High-content screening for neurite morphology analysis

Nuclei and neurites of neurons were imaged with Opera LX high-content screening system (PerkinElmer, Waltham, MA). The 96-well plates were scanned with 10x magnification, sampling 40 images per well. All images were analyzed using Columbus software (PerkinElmer, v2.5.2). Briefly, neuron nuclei were dissociated from non-neuronal nuclei or debris based on morphological (nuclei area > 200 µm^2^) and intensity (nuclei intensity > 100) parameters. Analyses were made at the population level of a well, taking the mean of selected neurons. Total neurite length (traced from the soma), number of protrusions (neurites branching) from the soma, number of first bifurcations (from soma protrusions), and number of extremities (neurite branching number from the first bifurcations) were calculated (see Fig. S1).

### Quantitative real time PCR for gene expression analysis

Transduced neurons were collected with TRIZOL reagent (Life Technologies) at DIV14 and RNA isolation was performed accordingly using isopropanol (Sigma Aldrich) precipitation (Spijker et al., 2004). RNA was quantified (NanoDrop ND-1000 spectrophotometer; NanoDrop Technologies, Wilmington, DE). Random-primed (25 pmol; Eurofins MWG Operon; Ebersberg, Germany) cDNA synthesis was performed on individual RNA samples (100–125 ng total RNA) using MMLV reverse transcriptase (Promega). Real-time qPCR reactions (10 μL; Light cycler 480, Roche) and relative gene expression calculation were performed as described previously (Schmitz et al., 2017). All qRT-PCR primers were designed using Primer3.0 software and are listed (forward and reverse) as follows: *Gpr158*: 5′- AACACAGCCTAGATCCAGAAGAC-3′ and 5′- GGGTTGTTTGTGATCATCTTTTTA-3′; *Gapdh*: 5′- TGCACCACCAACTGCTTAGC-3′ and 5′- GGCATGGACTGTGGTCATGA-3′; *β-Actin*: 5′- GCTCCTCCTGAGCGCAAG-3′ and 5′- CATCTGCTGGAAGGTGGACA-3′; *Hprt*: 5′- ATGGGAGGCCATCACATTGT-3′ and 5′- ATGTAATCCAGCAGGTCAGCAA-3′.

### Slice Preparation

All procedures were conducted in accordance with the Dutch license procedures and with approval by the VU Animal Experimentation Ethics Committee. Upon completion of the behavioural assessment, mice were swiftly decapitated and brains were extracted and placed for a short period, during transport to slicing room, in carbogenated (95% O_2_, 5% CO_2_) iced-cold ‘slicing buffer’ containing (in mM), 70 NaCl, 2.5 KCl, 1.25 NaH_2_PO_4_.H_2_O, 5 MgSO_4_.7H_2_O, 1 CaCl_2_.2H_2_O, 70 Sucrose, 25 D-Glucose, 25 NaHCO_3_, 1 Na-Ascorbate, 3 Na-Pyruvate; pH 7.4, 305mOsm. Horizontal hippocampal slices were retrieved at 300 µm thickness, using a vibrating-blade microtome (HM-650V, Thermo Scientific), in ice-cold carbogenated ‘slicing buffer’. Each slice was briefly washed in ‘holding ACSF’ and placed in a slices chamber filled with carbogenated ‘holding ACSF’ containing in mM, 125 NaCl, 3 KCl, 1.25 NaH_2_PO_4_.H_2_O, 2 MgCl_2_.6H_2_O, 1.3 CaCl_2_.2H_2_O, 25 D-Glucose, 25 NaHCO_3_, 1 Na-Ascorbate, 3 Na-Pyruvate; pH 7.4, 305 mOsm. Slices were left to recover at RT for at least one hour before recordings, and for the duration of the experimental day slices were maintained in the slice chamber containing ‘holding ACSF’.

### Spontaneous Excitatory and Inhibitory Postsynaptic Current Recordings

Individual slices were transferred to a submerged recording chamber and left to equilibrate for 10 minutes, under continuous perfusion, at ∼2 mL/min, with carbogenated ‘recording ACSF’ containing in mM, 125 NaCl, 3 KCl, 1.25 NaH_2_PO_4_.H_2_O, 1 MgCl_2_.6H_2_O, 1.3 CaCl_2_.2H_2_O, 25 D-Glucose, 25 NaHCO_3_; pH 7.4, 305 mOsm. Hippocampal CA1 pyramidal neurons were visualized under differential interference contrast microscopy, and selected based on their morphology. For sEPSC recordings in WT 10 cells from 4 animals, and in KO 14 cells from 4 animals were analyzed. For sIPSC recordings in WT 11 cells from 4 animals, and in KO 14 cells from 4 animals were analyzed. Whole-cell patch-clamp configuration was achieved using standard borosilicate glass pipettes, ∼3-5 Mohm, filled with an Caesium-gluconate based intracellular containing in mM, 130 Cs-gluconate, 8 NaCl, 10 HEPES, 0.3 EGTA, 4 ATP-Mg, 10 K_2_-Phosphocreatine, 0.3 GTP, 3 QX314-Cl, 0.3% Biocytin; pH 7.3, 290 mOsm. Recordings were conducted at ∼32 °C. Upon achieving a stable whole-cell configuration, up to seven minutes of sEPSCs were recorded at a holding potential of -70 mV, and up to seven minutes of sIPSCs were recorded at a holding potential of 0 mV. The last three minutes of each trace were analyzed with mini Analysis software (Synaptosoft). Acquisition was performed using p-Clamp software (Molecular Devices), using a Multiclamp 700B amplifier (Molecular Devices), sampled at 20 kHz, low-pass filtered at 6 kHz, and digitized with an Axon Digidata 1440A (Molecular Devices). Series resistance was monitored and only cells exhibiting less than 20% change were used for analysis. Cells whose access resistance exceeded 25 Mohm were also rejected.

### Paired Pulse Ratio Recordings

For PPR recordings whole-cell patch-clamp configuration was achieved with a K-gluconate based intracellular containing in mM, 148 K-gluconate, 1 KCl, 10 HEPES, 0.3 EGTA, 4 ATP-Mg, 4 K_2_-Phosphocreatine, 0.4 GTP, 0.3% Biocytin; pH 7.3, 290 mOsm. For this experiment in WT 10 cells from 4 animals, and in KO 10 cells from 3 animals were analyzed. Upon achieving stable whole-cell configuration action potential profiles were generated for each CA1 pyramidal, through current injections of 750 ms starting from -200 pA, at steps of 20 pA. Subsequently, a unipolar stimulating electrode loaded with ‘recording ACSF’ was lowered near the location were Schaffer collaterals meet the patched-cell’s dendrite. The electrode was moved around until a clear unisynaptic response could be observed, upon current injections mediated by a Master-9 pulse stimulator and an ISO-flex stimulus isolator (A.M.P.I). Thereafter, an input-output stimulation curve was generated by recording the patched cells’ response to increasing amounts of current injections from 20 µA to 200 µA, at steps of 20 µA. For each step, 5 sweeps were averaged. The current intensity used during the PPR protocol was approximately the current generating half-maximal amplitude responses. The two pulses for the PPR protocol were separated by 50 ms, and 20-25 sweeps were recorded per cell. Data were analysed with in-house Matlab (Mathworks) scripts. All other electrophysiological and technical parameters were as described in the section regarding sEPSC/sIPSC recordings.

### Biocytin Reconstructions

Upon completion of an electrophysiological recordings the pipette was slowly retracted to reseal the patched cell, and the slice was subsequently fixed in 4% paraformaldehyde (PFA), at 4 °C for 48 h. PFA was replaced with PBS-Azide for long-term storage at 4 °C until further analysis. The following set of cells was reconstructed. From the cells recorded under the PPR section, in WT 10 cells from 4 animals, and in KO 10 cells from 3 animals were analyzed. From the cells under the sEPSC/sIPSC section, in WT 8 cells form 3 animals (one animal is shared with the PPR experiment), and in KO 7 cells from 3 animals. For the per animal morphology, the aforementioned cells were averaged per animal, yielding in WT 18 cells from six animals, and in KO 17 cells from 6 animals. Slices containing biocytin filled cells, were incubated in 3% H_2_O_2_ in 0.05 M PB solution for 20 min, and then incubated for 48 h in 0.05 M PB containing 0.5% Triton X-100, and ABC complex solution (Vectastain; 1 drop solution A, 1 drop solution B, per 20 mL volume). Subsequently, slices were incubated in 0.05 M PB containing 3.3 mM H_2_O_2_ and 2 mM diaminobenzidine tetrahydrochloride (DAB) under visual inspection. Incubation was terminated when a clear morphology could be visualized. All steps were separated by 3x washes in 0.05 M PB. Images of labelled cells were acquired with a light microscope (Olympus) at 20X magnification, using the Surveyor software (Objective Imaging). Reconstruction of the imaged cells was performed using the Neuromantic software (Myatt et al., 2012), and analysis of the reconstructed morphologies was performed using L-Measure (Scorcioni et al., 2008).

### Statistics

For genotype comparisons, two-tailed Student’s t-tests (with or without correction for unequal variation) were applied for normally distributed data and Mann-Whitney U-tests otherwise. Normality was assessed with Kolmogorov-Smirnov and Saphiro-Wilk tests. In the case of normally distributed data but with unequal variances, Welch’s unequal variance t-test was applied. Mixed ANOVA tests were carried out for genotype as between-subject factor and time as repeated measures (e.g., MWM training, PA extinction, I/O curve). A 2-way ANOVA was applied to the morphological analyses at the animal level (Fig. 5a) with genotype and type of dendrite as between-subject factors. All data pertaining to Principal component analysis (PCA) Figure S3 were generated using Matlab scripts (Mathworks). Multiple comparison correction for the correlation between cell surface and PCA variables was performed with the Bonferroni-Holm method. Two mice were removed from the MWM data set as they were floating instead of swimming (*Gpr158* KO n=1, WT n=1). For electrophysiology, cells with values exceeding mean±2xSD were excluded (PPR/Properties: *Gpr158* KO n=1, WT=3; sEPSC/sIPSC: *Gpr158* KO n=4, WT n=3). Statistical significance level was set for *P*-values<0.05. Statistical significance was assessed using SPSS v24 IBM, or Graphpad Prism 5 software (GraphPad Software, La Jolla, CA) for Bonferroni post-hoc tests.

## Supporting information

Supplements to main figures

## Acknowledgements

We thank Joost Hoetjes, Frank den Oudsten, Robert Zalm, Marion Sassen for their technical assistance. Additionally, we would like to thank Dr. Christian de Kock for access to the morphology acquisition recourses and Dr. Oliver Stiedl for setting up the passive avoidance test.

## Funding sources

DC was supported by CognitionNet EU-ITN MEST-CT-2013-607508. IK was supported by the NWO-TOP grant (91215030); ABS, HDM, SS received support from HEALTH-2009-2.1.2-1 EU-FP7 ‘SynSys’ (#242167); ABS, and RJvdL by NBSIK PharmaPhenomics grant LSH framework FES0908; HDM by an NWO VICI grant (ALW-Vici 865.13.002) and the ERC grant BrainSignals (281443); IK and SS by an NWO VICI grant (ALW-Vici 016.150.673/865.14.002).

## Competing interest

ABS participates in a holding that owns shares of Sylics BV. The authors declare no competing financial interests derived from this paper.

