## Supplements to main figures for "*Gpr158* deficiency impacts hippocampal CA1 neuronal excitability, dendritic architecture, and affects spatial learning"

**Supplemental files to the figures and main text**
**of**

***Gpr158* deficiency impacts hippocampal CA1 neuronal excitability, dendritic architecture, and affects spatial learning**

Demirhan Çetereisi*[,1], Ioannis Kramvis*[,1], Titia Gebuis[1], Rolinka J. van der Loo[1], Yvonne Gouwenberg[1], Huibert D. Mansvelder[2], Ka Wan Li[1], August B Smit[1], Sabine Spijker[1]

Departments of [1]Molecular and Cellular Neurobiology, and [2]Integrative Neurophysiology, Center for Neurogenomics and Cognitive Research, Amsterdam Neuroscience, Vrije Universiteit, Amsterdam, The Netherlands
* Equal contribution

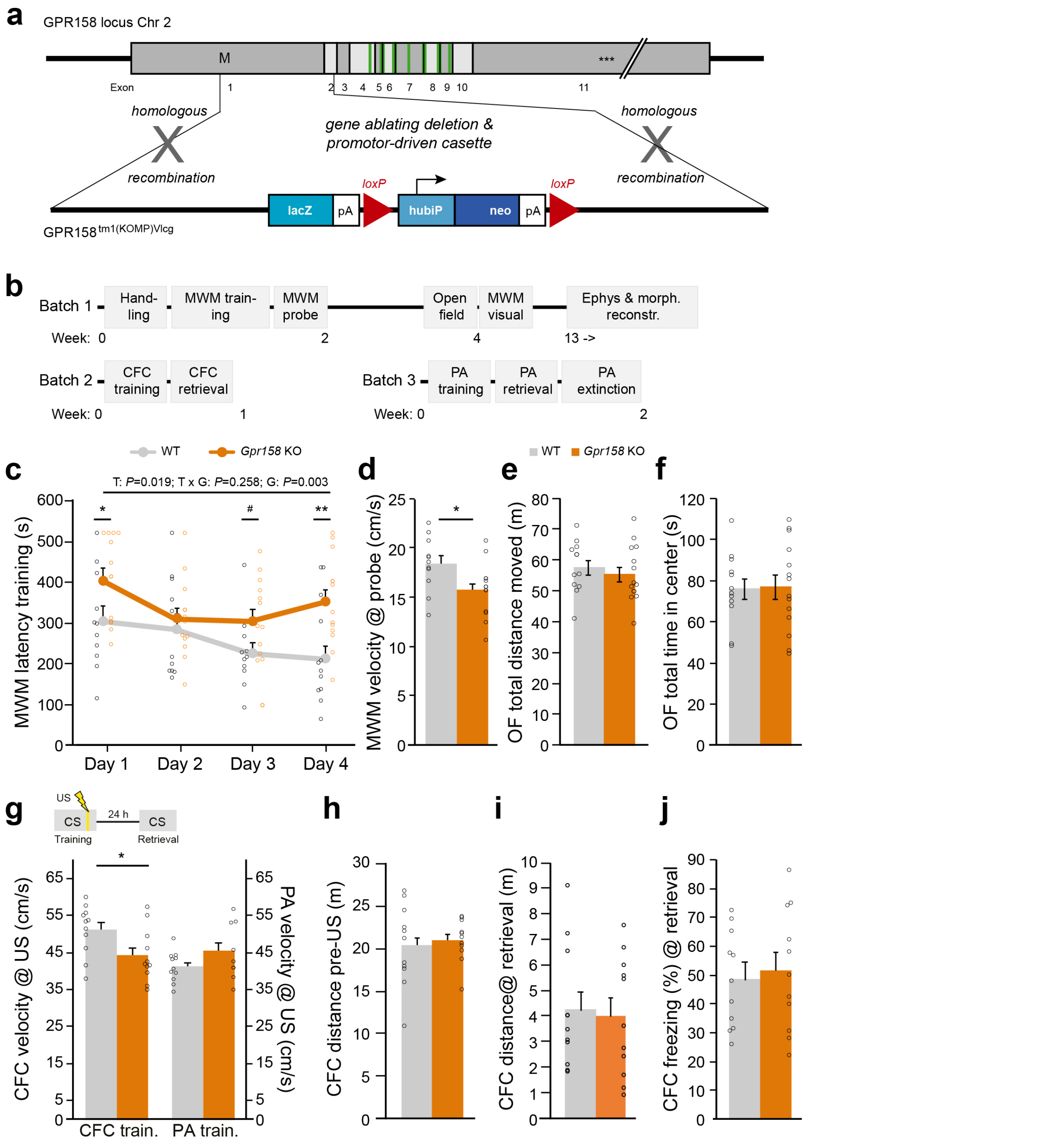


Figure 1 supplement 1. ***Gpr158* KO mice show a MWM acquisition deficit but normal open field and contextual fear memory**. **a**) Depicted is the exon structure of Gpr158 with 11 exons, with the approximate start site (M), stop site (***) and transmembrane regions (green bars). Schematic representation of the generation of *Gpr158* KO mice with the *Gpr158* locus and the Gpr158 KOMP construct (VG10108) used for homologous recombination (Gpr158$^{tm1(KOMP)Vlcg}$), generated previously (Orlandi et al., 2015), in which half of exon 1 and exon 2 were replaced by a LacZ cassette. **b**) Overview of the 3 batches used for behavioral analysis. **c,d**) MWM latency to reach the platform during training (c) showed an overall similar effect as distance (see Fig. 1a) with a genotype (G, *P*=0.003) and training (T, *P*=0.019) effect. *Gpr158* KO mice showed a significant difference in the mean velocity during the probe test (d), arguing that distance to find the platform during acquisition best reflects true genotype differences. **e,f**) Open field (OF) total distance moved (e; *P*=0.509), and total time spent in the center (f; *P*=0.909) showed no genotype differences (Figure 1 supplement 2; *Gpr158* KO n=14, WT n=12). **g-j**) Experimental set-up and data for testing long-term contextual fear memory (CFC). Mice received a foot shock (unconditioned stimulus, US) in the training context (conditioned stimulus, CS). Long-term memory was measured 24 h after training by placing the mouse back in the CS and measuring freezing level. *Gpr158* KO mice showed a small decrease in velocity after delivery of the foot shock in the CFC (g, left axis; *P*=0.031), possibly indicating a difference in shock perception. However, as this was not observed with the same shock intensity in the PA test in an independent batch (g, right axis; *P*=0.172, see Fig 1c), this is most likely a batch effect. *Gpr158* KO mice did not show a difference in terms of distance moved either during training prior to US delivery (h, *P*=0.715), nor 24 h later at memory retrieval (i, *P*=0.829). *Gpr158* KO mice did not show an impaired long-term memory in terms of their freezing level (j; *P*=0.693) upon memory retrieval, corroborating the intact PA memory (see Figure 1c). Previously a robust anxiety phenotype was detected (Khrimian et al., 2017). Here, we do not report such anxiety-like phenotype, neither in a novel open field, nor when placed in the fear conditioning box prior to conditioning. This might relate to the fact that the genetic background of the mice of the Knockout Mouse Project (KOMP) repository is C57BL/6NTac and Khrimian *et al.* kept their mice on a mixed 129-Sv/C57BL/6J background (Khrimian et al., 2017). The 129-Sv background is known for its anxiety (Rodgers et al., 2002; Tarantino et al., 2000), even when used in a mixed background (Võikar et al., 2001), and anxiety can be a confounding factor on cognitive performance (Brooks et al., 2005). Supporting this explanation is that *Gpr158* KO mice kept on a Bl6J background even showed reduced anxiety in the elevated plus maze (Sutton et al., 2018). Data are presented as mean±SEM with individual data points indicated. Asterisks and octothorpe indicate the level of significance between WT and KO assessed by two-tailed Student's t-test or MWU (Figure 1 supplement 2). # *P*≤0.10; * *P*≤0.050; ** *P*≤0.010.

| **Figure reference** | **n-number** | **type of overall or post-hoc test** | **(Df) F- or t-value** | ***P*-value** | **Remarks** |
|---|---|---|---|---|---|
| **1a**: MWM training distance | KO: 13 (& 1 floater taken out)<br>WT: 11 (& 1 floater taken out) | • mixed ANOVA: repeated measure (time) ANOVA (genotype)<br>• ttest | • time: (3,66) 0.80; time x genotype: (3,66) 2.61; genotype: (1,22) 4.49<br>• day 1: (22) -1.29; day 2: (22) 0.53; day 3: (22) -2.33; day 4: (22) -2.41 | • time: **<0.001**; time x genotype: **0.008**; genotype: **0.033**<br>• day 1: 0.209; day 2: 0.600; day 3: **0.031**; day 4: **0.025** | Spericity assumed (ANOVA); day 3: unequal variance |
| **1b**: MWM probe test time spent | KO: 11<br>WT: 14 | • mixed ANOVA: repeated measure (quadrant) ANOVA (genotype)<br>• ttest | • quadrant x genotype: (3,66) 4.46; quadrant for WT: (3,30) 3.45; quadrant for KO: (3,36) 2.23<br>• TQ: (22) 2.88; LQ: (22) 0.88; RQ: (19.5) -1.43; OQ: (22) -2.26 | • quadrant x genotype: **0.007**; quadrant for WT: **0.029**; quadrant for KO: 0.102<br>• TQ: **0.009**; LQ: 0.391; RQ 0.168; OQ **0.034** | Spericity assumed (ANOVA); RQ: unequal variance |
| **1c**: PA retrieval latency to enter | KO: 9<br>WT: 11 | MWU | | 0.656 (MWU) | non-normal |
| **1d**: PA extinction latency to enter | KO: 9<br>WT: 11 | • mixed ANOVA: repeated measure (time) ANOVA (genotype)<br>• ttest | • time: (2.34,42.20) 20.72; time x genotype: (2.34,42.20) 2.82; genotype: (1,18) 0.025<br>• day 1: (11.45) -2.14; day 2: (10.05) -2.37; day 3: (10.33) -1.97; day 4: (9.69) -1.74 | • time: **<0.001**; time x genotype: 0.065; genotype: 0.914<br>• day 1: 0.054; day 2: **0.039**; day 3: 0.077; day 4: 0.114 | Huyn-Feldt (ANOVA); day 1-4: unequal variance |
| **S1c**: MWM training latency | KO: 13<br>WT: 11 | • mixed ANOVA: repeated measure (time) ANOVA (genotype)<br>• ttest or MWU | • time: (3,66) 3.55; time x genotype: (3,66) 1.38; genotype: (1,22) 11.50<br>• day 3: (22) -1.91; day 4 (22) -3.06 | • time: **0.019**; time x genotype: 0.258; genotype: **0.003**<br>• day 1: **0.041** (MWU) ; day 2: 0.494 (MWU); day 3: 0.069 (ttest; day 4: **0.006** (ttest) | Spericity assumed (ANOVA); day 1 & 2: non-normal |
| **1S1d**: MWM distance probe test | KO: 13<br>WT: 11 | ttest | (22) 2.39 | **0.026** | |
| **1S1e**: OF total distance moved (m) | KO: 14<br>WT: 12 | ttest | (24) 0.67 | 0.509 | |
| **1S1f**: OF time spent in center (s) | KO: 14<br>WT: 12 | ttest | (24) -0.12 | 0.909 | |
| **1S1g**: cFC training, Vmean at US | KO: 11<br>WT: 11 | ttest | (20) 2.33 | **0.031** | |
| **1S1g**: PA Vmean at US | KO: 9<br>WT: 11 | ttest | (11.79) -1.54 | 0.172 | unequal variance |
| **1S1h**: cFC training, distance pre-US (m) | KO: 11<br>WT: 11 | ttest | (15.09) -0.37 | 0.715 | unequal variance |
| **1S1i**: cFC training, distance @ retrieval (m) | KO: 11<br>WT: 11 | ttest | (20) 0.22 | 0.829 | |
| **1S1j**: cFC retrieval, % freezing | KO: 11<br>WT: 11 | ttest | (20) -0.40 | 0.693 | |
| Main text: Visual test latency | KO: 13<br>WT: 11 | ttest | (18.10) -1.33 | 0.201 | unequal variance |

Figure 1 supplement 2. **Overview statistics behavioral assessments**. Shown are the statistical analyses (n-number, type of test, Df/F/t-value, *P*-value) for the data shown in Figure 1, and Figure 1 supplement 1 (1S1) related to behavior in *Gpr158* KO and WT animals. Significance (*P*<0.050) is indicated in bold, trend (*P*<0.100) in underlined.

| **Figure reference** | **n-number** | **type of overall or post-hoc test** | **(Df) & F- or t-value** | ***P*-value** | **Remarks** |
|---|---|---|---|---|---|
| **2b**: Frequency sEPSCs (Hz) | KO: 14 (4 animals)<br>WT: 10 (4 animals) | ttest | (22) 1.34 | 0.193 | unequal variance |
| **2b**: Current sEPSCs (pA) | KO: 14 / 4<br>WT: 10 / 4 | MWU | | 0.509 | non-normal |
| **2b**: Decay time sEPSCs (ms) | KO: 14 / 4<br>WT: 10 / 4 | ttest | (20.3) -2.12 | **0.047** | unequal variance |
| **2b**: Rise time sEPSCs (ms) | KO: 14 / 4<br>WT: 10 / 4 | ttest | (20.3) -1.39 | 0.179 | |
| **2d**: Frequency sIPSCs (Hz) | KO: 14 / 4<br>WT: 11 / 4 | ttest | (23) 0.734 | 0.470 | |
| **2d**: Current sIPSCs (pA) | KO: 14 / 4<br>WT: 11 / 4 | MWU | - | - | non-normal |
| **2d**: Decay time sIPSCs (ms) | KO: 14 / 4<br>WT: 11 / 4 | ttest | (23) -0.01 | 0.995 | |
| **2d**: Rise time sIPSCs (ms) | KO: 14 / 4<br>WT: 11 / 4 | ttest | (23) -1.03 | 0.314 | |
| **2f**: input-output | KO: 10 / 3<br>WT: 10 / 4 | • mixed ANOVA: repeated measure (stimulation) ANOVA (genotype)<br>• MWU | • stimulation: (1.34,24.13) 41.4; time x genotype: (1.34,24.13) 9.36; genotype: (1,18) 8.37 | • stimulation: **<0.0001**; stimulation x genotype: **0.003**; genotype: **0.001**<br>• 20 ms: 0.315; 40 ms: **0.023**; 60 ms: **0.003**; 80 ms: **0.003**; 100 ms: **<0.001**; 120 ms: **0.001**; 140 ms: **0.001**; 160 ms: **0.002**; 180 ms: **<0.001**; 200 ms: **0.001** | • Greenhouse-Geisser<br>• All non-normal |
| **2g**: Paired-pulse ratio | KO: 10 / 3<br>WT: 10 / 4 | ttest | (12.8) 0.06 | 0.954 | unequal variance |
| **2h**: Current (pA) by pulse | KO: 10 / 3<br>WT: 10 / 4 | ttest | 1st pulse: (18) 5.01<br>2nd pulse: (18) 5.70 | 1st pulse: **<0.001**<br>2nd pulse: **<0.001** | |

Figure 2 supplement 1. **Overview statistics spontaneous EPSCs / IPSCs, and evoked responses**. Shown are the statistical analyses (n-number (slice / animal), type of test, Df/F/t-value, *P*-value) for the data shown in Figure 2, related to electrophysiological assessment in *Gpr158* KO and WT CA1. Significance (*P*<0.050) is indicated in bold.

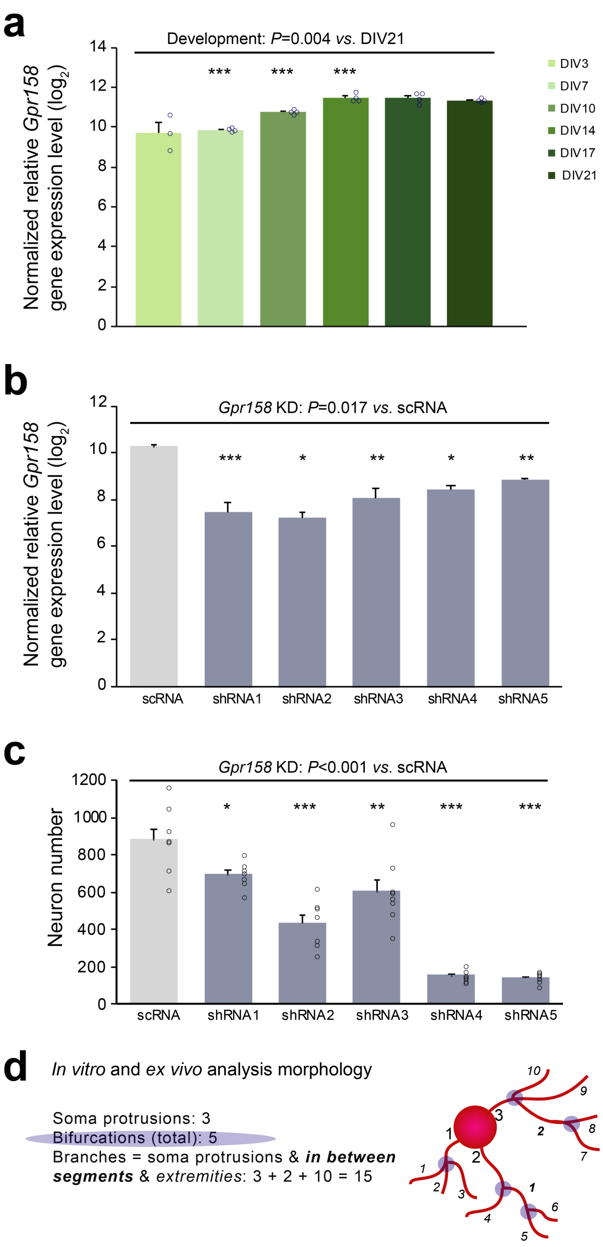


Figure 3 supplement 1. ***Gpr158* knock-down (KD) in hippocampal culture**. **a**) Endogenous gene expression level of *Gpr158* from an *in vitro* hippocampal primary culture (n=4 wells, except for DIV3, n=3 wells) during development (DIV3-21). *Gpr158* showed an overall developmental increase in gene expression levels (Kruskal-Wallis, *P*=0.004), with expression being significantly different from DIV7 to DIV14 as compared to DIV21 (Figure 3 supplement 2). **b**) Knock-down (KD) efficiency of five shRNAs against the *Gpr158* gene. Neurons were transduced at DIV7 and harvested at DIV14 Overall, all shRNAs significantly reduced *Gpr158* gene expression level compared with the scrambled control (scRNA) (Kruskal-Wallis, *P*=0.017, Figure 3 supplement 2). **c**) *Gpr158* KD effect on neuron number in hippocampal primary cultures. Neurons were transduced at DIV7 and the neuron number was counted at DIV14. It is of note that the large reduction in neuronal number in shRNA4 and 5 might have left a higher proportion of non-transduced neurons alive, resulting in more variable results with respect to endogenous *Gpr158* expression (see panel b), or to specific morphological parameters (see Figure 3). **d**) Schematic representation of morphological parameters analyzed for *in vitro* (see Figure 3) and *ex vivo* (see Figure 4). Purple shaded circles indicates the number of bifurcations. Data are presented as mean±SEM with individual data points indicated. Asterisks indicate the level of significance between WT and KO assessed by Student's t-test (Figure 3 supplement 2). * $P\leq0.050$; ** $P\leq0.010$; *** $P\leq0.001$.

| **Figure reference** | **n-number** | **type of overall or post-hoc test** | **(Df) & F- or t-value** | ***P*-value** | **Remarks** |
|---|---|---|---|---|---|
| **3c**: Total neurite length | Sc: 8 wells<br>shRNA1: 7 wells;<br>shRNA2: 7 wells;<br>shRNA3: 8 wells;<br>shRNA4: 7 wells;<br>shRNA5: 8 wells | • Kruskal-Wallis (all shRNAs *vs.* Sc)<br>• ttest *vs.* Sc | • -<br>• shRNA1: (13) -0.16; shRNA2: (13) 0.83; shRNA3: (14) 7.53; shRNA4: (8.33) 1.73; shRNA5: (14) 8.58 | • treatment: <**0.001**<br>• shRNA1: 0.870; shRNA2: 0.424; shRNA3: **<0.001**; shRNA4: 0.120; shRNA5: **<0.001** | • Unequal variance in ANOVA<br>• shRNA4, unequal variance |
| **3d**: Soma protrusions | See above | • Kruskal-Wallis (all shRNAs *vs.* Sc)<br>• ttest *vs.* Sc | • -<br>• shRNA1: (8.75) -3.82; shRNA2: (13) -3.04; shRNA3: (14) 2.84; shRNA4: (13) 11.60; shRNA5: - | • treatment: **<0.001**<br>• shRNA1: **0.004**; shRNA2**: 0.010**; shRNA3: **0.013**; shRNA4: **<0.001**; shRNA5: 0.234 (MWU) | • Unequal variance in ANOVA<br>• shRNA1, unequal variance; shRNA 5, non-normal |
| **3e**: Bifurcation | See above | • Kruskal-Wallis (all shRNAs *vs.* Sc)<br>• ttest *vs.* Sc | • -<br>• shRNA1: (13) 0.735; shRNA2: (13) 2.297; shRNA3: (7.724) 7.600; shRNA4: (7.064) -0.614; shRNA5: (8.040) 8.433 | • treatment: **<0.001**<br>• shRNA1: 0.476; shRNA2: **0.039**; shRNA3: **<0.001**; shRNA4: 0.558; shRNA5: **<0.001** | • Unequal variance in ANOVA<br>• shRNA3-5, unequal variance |
| **3f**: Extremities | See above | • Kruskal-Wallis (all shRNAs *vs.* Sc)<br>• ttest *vs.* Sc | • -<br>• shRNA1: (13) 0.244; shRNA2: (13) 1.393; shRNA3: (14) 7.024; shRNA4: (8.149) 0.793; shRNA5: (14) 7.908 | • treatment: **<0.001**<br>• shRNA1: 0.811; shRNA2: 0.187; shRNA3: **<0.001**; shRNA4: 0.450; shRNA5: **<0.001** | • Unequal variance in ANOVA<br>• shRNA4, unequal variance |
| **3S1a**: Normalized relative *Gpr158* expression level (log2) | DIV3: 2 wells; rest: 4 wells | • Kruskal-Wallis (development)<br>• ttest *vs.* DIV21 | • -<br>• DIV3: (1.03) 5.04; DIV7: (6) 20.66; DIV10: (6) 7.03; DIV14: (6) 7.03; DIV17: (6) -0.86 | • development: **0.004**<br>• DIV3: 0.119; DIV7: **<0.001**; DIV10: **<0.001**; DIV14: **<0.001**; DIV17: 0.421 | • Unequal variance in ANOVA<br>• DIV3: unequal variance |
| **3S1b**: Normalized relative *Gpr158* expression level (log2) | shRNA5: 2 wells; rest: 3 wells | • Kruskal-Wallis (all shRNAs vs. Sc)<br>• ttest *vs.* Sc | • -<br>• shRNA1: (4) 19.44; shRNA2: (2.13) 6.47; shRNA3: (4) 9.08; shRNA4: (2.15) 4.12; shRNA5: (4) 8.15 | • treatment: **0.017**<br>• shRNA1: **<0.001**; shRNA2: **0.020**; shRNA3: **0.001**; shRNA4: **0.048**; shRNA5: **0.004** | • Unequal variance in ANOVA<br>• shRNA2 & shRNA4: unequal variance |
| **3S1c**: Neuron number | Sc: 8 wells<br>shRNA1: 7 wells;<br>shRNA2: 7 wells;<br>shRNA3: 8 wells;<br>shRNA4: 7 wells;<br>shRNA5: 8 wells | • Kruskal-Wallis (all shRNAs *vs.* Sc)<br>• ttest vs. Sc | • -<br>• shRNA1: (13) 2.70; shRNA2: (13) 5.61; shRNA3: (14) 3.15; shRNA4: (7.54) 11.74; shRNA5: (7.36) 11.95 | • treatment: **<0.001**<br>• shRNA1: **0.018**; shRNA2: **<0.001**; shRNA3: **0.007**; shRNA4: **<0.001**; shRNA5: **<0.001** | • Unequal variance in ANOVA<br>• shRNA4&5, unequal variance |

Figure 3 supplement 2**. Overview statistics *in vitro* morphological analysis**. Shown are the statistical analyses (n-number (wells), type of test, Df/F/t-value, *P*-value) for the data shown in Figure 3 and Figure 3 supplement 1 (3S1), related to the *in vitro* analysis of *Gpr158* expression and *Gpr158* KD in WT hippocampus primary neurons. Significance (*P*<0.050) is indicated in bold.

| **Figure reference** | **n-number** | **type of overall or post-hoc test** | **(Df) & F- or t-value** | ***P*-value** | **Remarks** |
|---|---|---|---|---|---|
| **4b**: Length (mm) | KO: 10 (3 animals); WT: 10 (4 animals) | ttest | Total: (18) 3.63<br>Apical: (18) 3.76<br>Basal: (18) 0.97 | Total: **0.002**<br>Apical: **0.001**<br>Basal: **0.346** | |
| **4b**: Surface area (mm2) | KO: 10 / 3 WT: 10 / 4 | ttest | Total: (18) 3.65<br>Apical: (18) 3.73<br>Basal: (18) 1.56 | Total: **0.002**<br>Apical: **0.002**<br>Basal: 0.136 | |
| **4b**: Bifurcations (#) | KO: 10 / 3 WT: 10 / 4 | ttest, MWU | (13.4) 3.76<br>Apical: -<br>Basal: - | Total: **0.002**<br>Apical: **0.001**<br>Basal: 0.481 | Total: unequal variance; Apical & basal: non-normal |
| **4b**: Branches (#) | KO: 10 / 3 WT: 10 / 4 | ttest, MWU | (14.16) 3.63<br>Apical: -<br>Basal: - | Total: **0.003**<br>Apical: **0.001**,<br>Basal: 0.529 | Total: unequal variance; Apical & basal: non-normal |
| **4d**: Amplitude (mV) | KO: 10 / 3 WT: 10 / 4 | ttest | (10.7) 0.80 | 0.432 | |
| **4d**: Threshold (mV) | KO: 10 / 3 WT: 10 / 4 | ttest | (12.1) -1.06 | 0.311 | |
| **4d**: Minimal ISI (ms) | KO: 10 / 3 WT: 10 / 4 | MWU | - | **0.015** | non-normal |
| **4d**: Rheobase (pA) | KO: 10 / 3 WT: 10 / 4 | MWU | - | **0.009** | non-normal |
| **4d**: Input resistance (MOhm) | KO: 10 / 3 WT: 10 / 4 | ttest | (18) -3.337 | **0.004** | |
| **4d**: Membrane potential (mV) | KO: 10 / 3 WT: 10 / 4 | MWU | - | **0.035** | non-normal |

Figure 4 supplement 1. **Overview statistics *ex vivo* morphological and AP profile analysis**. Shown are the statistical analyses (n-number (cells / animals), type of test, Df/F/t-value, *P*-value) for the data shown in Figure 4, related to the *ex vivo* morphological analysis of *Gpr158* KO and WT neurons in hippocampus CA1 that were analyzed for their intrinsic properties. Significance (*P*<0.050) is indicated in bold.

**a**

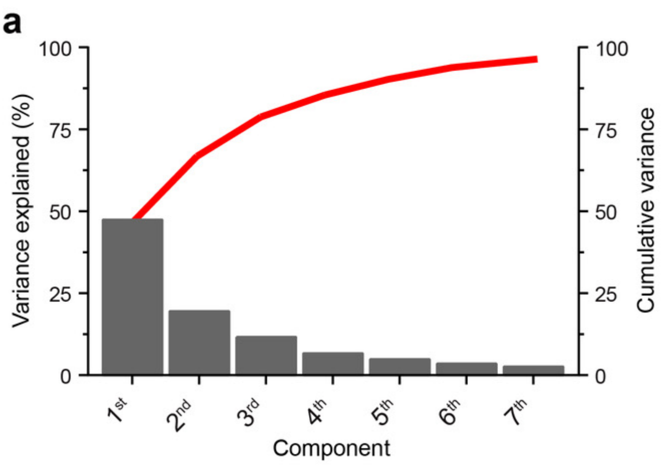


| | Variance explained (%) | Cumulative variance |
|---|---|---|
| **Component 1** | 47.4815623 | 47.481562 |
| **Component 2** | 19.5554565 | 67.037019 |
| **Component 3** | 11.7411817 | 78.778200 |
| **Component 4** | 6.67756245 | 85.455763 |
| **Component 5** | 4.85807496 | 90.313838 |
| **Component 6** | 3.52860925 | 93.842447 |
| **Component 7** | 2.63547906 | 96.477926 |
| **Component 8** | 1.84516526 | 98.323091 |
| **Component 9** | 0.70357794 | 99.026669 |
| **Component 10** | 0.51644760 | 99.543117 |
| **Component 11** | 0.21526819 | 99.758385 |
| **Component 12** | 0.16613684 | 99.924522 |
| **Component 13** | 0.06017517 | 99.984697 |
| **Component 14** | 0.01404516 | 99.998742 |
| **Component 15** | 0.00125764 | 100 |

**b**

| | Principal Component Analysis | | | Cell surface area *vs.* | | |
|---|---|---|---|---|---|---|
| | **1st Comp.** | **2nd Comp.** | **3rd Comp.** | **Pearson *r*** | ***P*-value** | ***P*-value corrected** |
| **Length** | 467.041 | 115.618 | 220.894 | 0.9789 | 0.0000 | **0.0000** |
| **Surface** | 6213.502 | 1940.175 | 2627.977 | 1.0000 | 0.0000 | **0.0000** |
| **Bifurcations** | 4.347 | 1.140 | 3.214 | 0.9348 | 0.0000 | **0.0000** |
| **Branches** | 8.891 | 2.386 | 6.747 | 0.9447 | 0.0000 | **0.0000** |
| **AP Amplitude** | -0.001 | -3.069 | 0,989 | -0.1324 | 0.5779 | - |
| **AP threshold** | 0.066 | 1.374 | -0.338 | 0.1518 | 0.5229 | - |
| **AP min ISI** | 1.809 | 1.140 | 0.428 | 0.7571 | 0.0001 | **0.0011** |
| **Rheobase free** | 8.080 | -1.496 | -3.452 | 0.6681 | 0.0013 | **0.0115** |
| **Vm** | -0.938 | 1.868 | -0.460 | -0.2261 | 0.3379 | - |
| **Ri** | -11.607 | 4.737 | -4.347 | -0.8458 | 0.0000 | **0.0000** |
| **TauM** | 0.743 | 0.813 | 0.878 | 0.3648 | 0.1138 | - |
| **PPR 1st Raw Amplitude** | 10.573 | -3.618 | -16.781 | 0.5277 | 0.0168 | 0.1343 |
| **PPR 2nd Raw Amplitude** | 16.449 | -11.538 | -22.662 | 0.4893 | 0.0286 | 0.1999 |
| **PPR** | -0.014 | -0.096 | 0.106 | -0.1428 | 0.5482 | - |
| **Stimulation Amplitude @ 200** | 25.717 | -25.706 | -59.982 | 0.3493 | 0.1312 | - |

Figure 4 supplement 2. **Principle component analysis of morphological and electrophysiological parameters**. **a**) Graph showing the variance explained by the first 7 PCA components (bars) and cumulative (red line). Table with all 15 components deduced from PCA of morphological and electrophysiological parameters measured from the same set of pyramidal neurons (see Figure 4). **b**) The coefficients for the different morphological and electrophysiological parameters of the first 3 components are indicated, demonstrating cell surface area as the most prominent determinant. To the right, the correlation matrix between cell surface area against all variables included in the PCA, including the Pearson correlation *r*, *P*-value, and multiple comparison corrected *P*-value (bold when significant).

| Figure reference | n-number | type of overall or post-hoc test | (Df) & F- or t-value | *P*-value | Remarks |
|---|---|---|---|---|---|
| **5a**: Length (mm) | KO: 6<br>WT: 6 | • 2-way ANOVA (compartment; genotype)<br>• Bonferroni correction (apical/basal) | • Compartment: (1,23) 219.10; genotype: (1,23) 13.39; interaction: (1,23) 5.22<br>• Apical: (20) 4.204<br>Basal: (20) 0.971 | • Compartment: **<0.001**; genotype: **0.002**; interaction: **0.033**<br>• Apical: **0.001**; Basal: 0.686 | |
| **5a**: Surface area (mm2) | KO: 6<br>WT: 6 | • 2-way ANOVA (compartment; genotype)<br>• Bonferroni correction (apical/basal) | • Compartment: (1,23) 199.64; genotype: (1,23) 17.73; interaction: (1,23) 4.91<br>• Apical: (20) 4.544<br>Basal: (20) 1.411 | • Compartment: **<0.001**; genotype: **0.001**; interaction: **0.001**<br>• Apical: **<0.001**; Basal: 0.347 | |
| **5a**: Bifurcations (#) | KO: 6<br>WT: 6 | • 2-way ANOVA (compartment; genotype)<br>• Bonferroni correction (apical/basal) | • Compartment: (1,23) 110.41; genotype: (1,23) 15.00; interaction: (1,23) 3.22<br>• Apical: (20) 4.008<br>Basal: (20) 1.470 | • Compartment: **<0.001**; genotype: **0.001**; interaction: <u>0.088</u><br>• Apical: **0.001**; Basal: 0.314 | |
| **5a**: Branches (#) | KO: 6<br>WT: 6 | • 2-way ANOVA (compartment; genotype)<br>• Bonferroni correction (apical/basal) | • Compartment: (1,23) 84.43; genotype: (1,23) 14.56; interaction: (1,23) 3.31<br>• Apical: (20) 3.984<br>Basal: (20) 0.347 | • Compartment: **<0.001**; genotype: **0.001**; interaction: <u>0.084</u><br>• Apical: **0.002**; Basal: 0.347 | |

Figure 5 supplement 1. **Overview statistics of morphological analysis for correlation with behavior**. Shown are the statistical analyses (n-number (animals), type of test, Df/F/t-value, *P*-value (ANOVA; Bonferroni correction) for data shown in Figure 5, related to the *Gpr158* KO and WT mice in which both morphological analysis of CA1 pyramidal neurons and Morris Water maze learning was performed. Significance (*P*<0.050) is indicated in bold, trend (*P*<0.100) in underlined.

| Name | symbol | # peptides | CA3 expression | CA1 expression | DG expression | Probe Allen Brain Atlas |
|---|---|---|---|---|---|---|
| Agrin | Agrn | 30 | high | high | medium | Probe RP_051214_03_B09 |
| Glypican1 | Gpc1 | 10 | high | medium | low | Probe RP_051214_03_E04 |
| Glypican2 | Gpc2 | 3 | - | - | - | |
| Glypican3 | Gpc3 | 7 | low | low | medium | Probe RP_050329_01_C10 |
| Glypican4 | Gpc4 | 6 | - | - | medium | Probe RP_040428_01_C03 |
| Glypican6 | Gpc6 | 8 | - | - | - | |
| Neuropilin1 | Nrp1 | 4 | high | low | medium | Probe RP_050915_01_H06 |
| Perlecan | Hspg2 | 33 | - | - | - | |
| Pikachurin | Egflam | 3 | - | - | - | |
| Syndecan4 | Sdc4 | 3 | - | - | - | |
| Spock1/Testican1 | Spock1 | 7 | high | low | medium | Probe RP_051101_02_A03 |
| Spock3/Testican3 | Spock3 | 11 | high | low | high | Probe RP_050725_01_F03 |

**Supplementary file: Overview of hippocampus subregion gene expression of ECM-related Gpr158 interactors**. For the published set of 12 ECM-related Gpr158 N-terminal interactors (Orlandi et al., 2018) the level of expression (high; medium; low; absent (-)) in CA3, CA1 and DG is indicated based on specific Allen Brain Atlas probes. Specifically, genes that are expressed in the CA3 region (yellow) are of interest, as they could subserve a similar role in the CA3 to CA1 pathway, as Glypican4 does in the MF to CA3 pathway, as elegantly shown before (Condomitti et al., 2018). It should be noted however that in total 129 Gpr158 N-terminal interactors were found (Orlandi et al., 2018), increasing the possibility of finding a similar functional pre-postsynaptic pair.
